# Stimulation of rodent and human beta-cell proliferation using synthetic modified mRNAs encoding cell cycle regulators

**DOI:** 10.64898/2026.08.21.746224

**Authors:** Tomas Koblas, Katerina Bittenglova, Pavel Abaffy, Klara Zacharovova, Peter Girman, Lukas Valihrach, Jan Kriz, Frantisek Saudek

## Abstract

Pancreatic beta cells exhibit marked resistance to proliferation, posing a barrier to therapeutic strategies aimed at restoring beta-cell mass in diabetes. Here, we present a transient, non-integrative approach to stimulate beta-cell proliferation using in vitro transcribed (IVT) mRNAs encoding cell cycle regulators. In rodent beta cells and human-beta cell derived EndoC-BH5 cells, chemically modified IVT mRNAs activated cell cycle entry and subsequent mitosis. A single dose of cyclin D1 and CDK4 IVT mRNAs nearly doubled the number of rat beta cells. However, achieving cell division in human beta cells required co-delivery of MYC IVT mRNA. The mitogenic response of beta cells peaked within 36–60 hours, and declined thereafter, reflecting the transient nature of IVT mRNA. Transcriptomic profiling revealed temporary activation of proliferative pathways and reversible downregulation of beta-cell maturation markers. Importantly, we detected no evidence of sustained proliferation. Our findings demonstrate that mRNA-based delivery of cell cycle regulators can overcome the intrinsic cell cycle block in beta cells and may provide a controllable approach for beta-cell regeneration.

## INTRODUCTION

Diabetes is a chronic metabolic disorder resulting from the loss or dysfunction of insulin-producing beta cells. Beta-cell loss is associated with the onset or progression of both type 1 and type 2 diabetes,^1,2^ ultimately resulting in a lifelong dependence on exogenous insulin supply and inadequate blood glucose control in people with diabetes.

Accordingly, restoring functional beta-cell mass is a major therapeutic goal in diabetes care. One promising therapeutic approach is the activation of beta-cell proliferation and subsequent increase in beta-cell number. However, except for a short period of proliferation during infancy, human beta cells show minimal physiological proliferation due to their resistance to cell cycle activation.^3^ Thus, only a few approaches have achieved a meaningful increase in human beta-cell proliferation. These include the use of small molecules that induce the expression of cell cycle regulators required for activation and completion of the cell cycle.^4–6^ Another approach involves using vectors to directly overexpress cell cycle regulators, such as cyclin-dependent kinases (CDKs) and their regulatory partners, cyclins.^7^ Although the viral-based approach significantly increases beta-cell proliferation, its therapeutic application is questionable. High viral doses can induce deleterious effects, such as inflammatory response, apoptosis, and a risk of insertional mutagenesis, limiting potential clinical application.^8,9^ Moreover, viral vectors often display unpredictable expression kinetics that do not reflect the tightly regulated expression profile of cell cycle genes^10,11^ and may compromise proper cell cycle progression.^12^

As an alternative to viral vectors, the genetic information encoding the protein of interest can be delivered to target cells using non-viral methods. These include DNA plasmids or synthetic *in vitro* transcribed messenger RNA (IVT mRNA).^13–15^ IVT mRNA structurally resembles natural mRNA; upon intracellular delivery, it allows transient overexpression of a specific protein without the risk of insertional mutagenesis.^15^ Unlike DNA vectors, IVT mRNA is translated directly in the cell cytoplasm, bypassing the need for nuclear entry and thus supporting high and rapid translation rates.^16^ Additionally, the highly regulated temporal lifespan of IVT mRNA enables transient, controllable expression of specific proteins.^17^ Nonetheless, IVT mRNA can be highly immunogenic and may induce an innate immune response in transfected cells, led by intrinsic antiviral defense mechanisms.^14,18^ Incorporating modified nucleosides into the IVT mRNA addresses this key issue and attenuates its immunogenicity.^19^ In particular, the substitution of unmodified uridine-5′-triphosphate with pseudouridine-5′-triphosphate or N1-methylpseudouridine triphosphate, and cytidine-5′-triphosphate with 5-methylcytidine-5′-triphosphate, decreases the inflammatory response.^20^ In addition, incorporating modified nucleosides improves the stability and translational efficiency of IVT mRNA.^21^ High-performance liquid chromatography or affinity purification can further improve the tolerability and translation efficiency of IVT mRNA.^22,23^ Both these approaches remove highly immunogenic double-stranded RNA contaminants—a by-product of IVT mRNA synthesis. The feasibility of IVT mRNA treatment has already been demonstrated in clinical trials for vaccines, cancer immunotherapy, gene editing, and protein replacement therapy.^14,15,17,24–27^ For instance, BioNTech/Pfizer and Moderna’s SARS-CoV-2 IVT mRNA vaccines were highly effective and safe in preventing symptomatic COVID-19 in Phase 3 clinical trials and received Conditional Marketing Authorizations or Emergency Use Authorizations in numerous countries, playing a key role in controlling the COVID-19 pandemic.^28,29^

In this study, we evaluated IVT mRNA technology for inducing beta-cell proliferation. Our findings indicate that IVT mRNA encoding specific cell cycle regulators can significantly increase proliferation in both rodent and human beta cells. While rodent beta-cell proliferation was induced by a simple combination of IVT mRNA for cyclins D and early CDKs, human beta cells required a more complex combination that included MYC IVT mRNA. Additionally, we confirmed effective beta-cell proliferation using two independent assays that assess not only cell cycle entry and progression but also mitotic cell division. This allowed us to determine up to a 2-fold increase in total rodent beta-cell mass following IVT mRNA-induced proliferation. We also verified that the stimulatory effect of IVT mRNA is temporary, with induced beta-cell proliferation ceasing within 3–4 days post-transfection.

## RESULTS

### IVT mRNA allows ectopic overexpression of cell cycle regulators in pancreatic beta cells

Cyclins and CDKs are key regulators that activate cell cycle entry and progression. Therefore, we first selected a group of cyclins and early CDKs previously shown to induce rodent and human beta-cell proliferation using viral vectors.^7^ Specifically, we designed and prepared IVT mRNAs encoding CDK4, CDK6, and cyclins D1, D2, D3. Additionally, we prepared mutant versions of cyclins D by replacing threonine for alanine at specific positions (cyclin D1 T286A; cyclin D2 T280A; cyclin D3 T283A). These mutations enhance the stability of the translated proteins, facilitate their preferential localization to the nucleus, and, as a result, increase the proliferation rate of expressing cells.^30^ Furthermore, all IVT mRNAs contained 5′ and 3′ untranslated regions (UTRs) derived from beta-cell specific insulin mRNA (*Ins2*) to improve IVT mRNA translation efficiency, especially in beta cells.

Next, we evaluated the efficiency of IVT mRNA in inducing expression of the encoded cyclins and CDKs upon transfection into rat pancreatic islet cells. Following transfection of islet cells with specific IVT mRNAs, we detected a significant overexpression of all encoded proteins within 16 hours post-transfection (hpt), as detected by immunofluorescence staining (IF) and Western blotting (WB) (Figures 1B and 1D). However, we did not find substantial differences in protein expression between wild-type (WT) and mutant forms of cyclins D. In addition, we did not detect notable expression of any cyclin D or CDK in untransfected control samples, except for cyclin D2 (Figure 1D). This result is in agreement with a previous report that revealed significant cyclin D2 expression in comparison with the other D-type cyclins and its essential role in postnatal beta-cell proliferation and function.^31^

**Figure 1.**
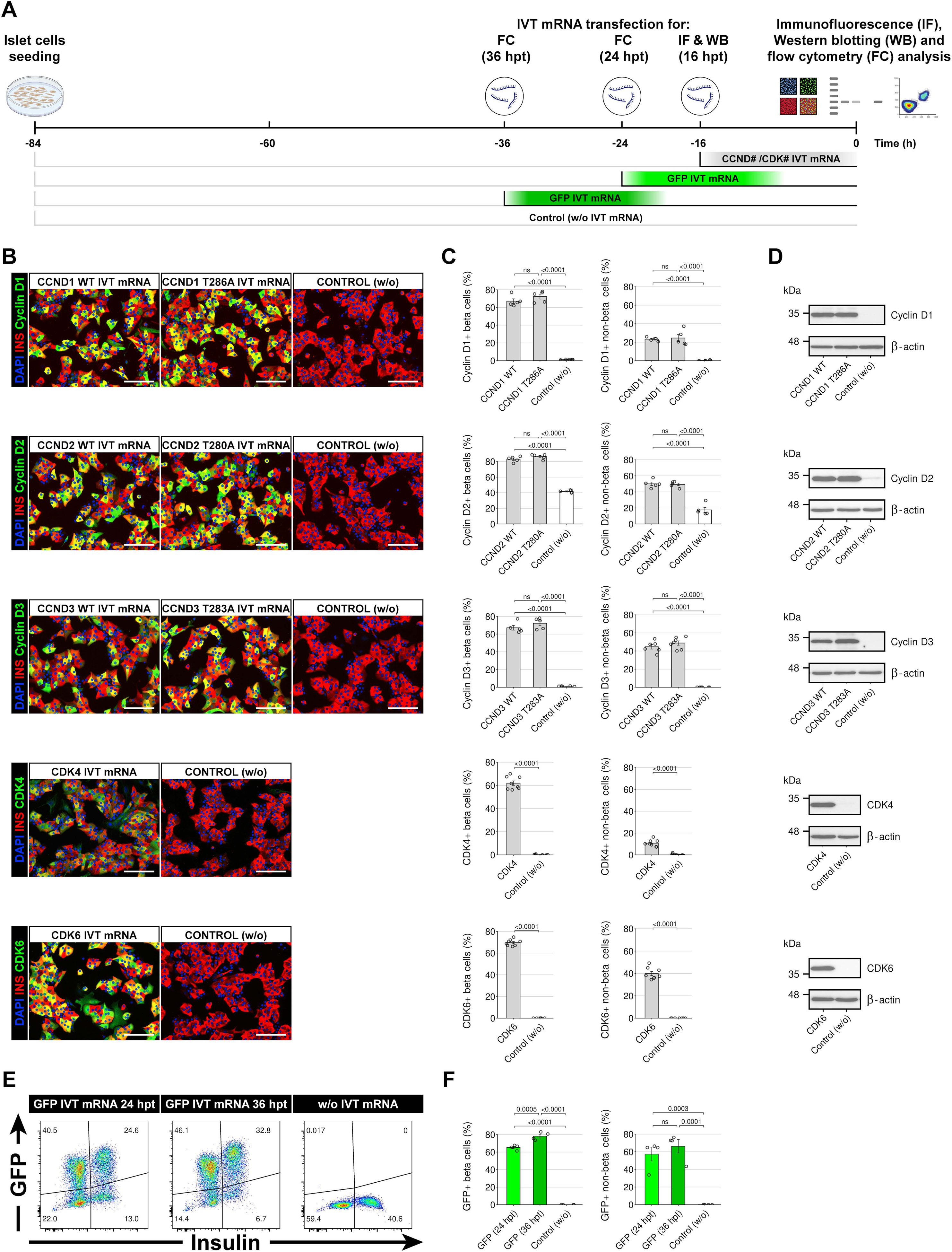
Ectopic Cyclin D/CDK overexpression in islet cells upon transfection with IVT mRNAs encoding specific cell cycle regulators. (A) Schematic showing the experimental design and timeline for testing cyclins D (CCND), CDK, and GFP IVT mRNA translation and transfection efficiency in islet cells. The figure was created using BioRender.com. (B) Representative merged images of islet cells at 16 h post-transfection (hpt) with IVT mRNA encoding specific cell cycle regulators (cyclins D/CDKs). Co-immunostaining for insulin (red) and cyclin/CDK (green). Nuclei are marked with DAPI (blue). Yellow, orange, and light green cells indicate beta cells co-expressing specific cyclin D/CDK according to their expression level. Untransfected islet cells were used as negative control. Scale bars, 100 μm. (C) Quantification of beta and non-beta-cell positivity for specific cyclin D/CDK after IVT mRNA transfection. Data represent mean ± standard deviation (SD) (n = 5 biologically independent experiments). Statistical significance was determined using one-way analysis of variance (ANOVA) with Bonferroni’s post-test analysis, or unpaired two-tailed Student’s t-test. P-values are shown above the graphs. (D) Western blot detection of cyclin D/CDK protein expression in islet cells at 16 hpt with IVT mRNA encoding specific proteins. Beta-actin was used as loading control. Note that cyclin D2 is detectable in untransfected control samples. (E) Flow cytometry analysis of GFP IVT mRNA transfection efficiency, at 24 and 36 h after transfection. (F) Quantification of GFP expressing beta and non-beta-cells after GFP IVT mRNA transfection. Data represent mean ± SD (n = 4). Statistical significance was determined using one-way ANOVA with Bonferroni’s post-test analysis. P-values are shown above the graphs.

Co-staining for insulin and specific cyclin D (CCND) or CDK revealed highly variable rates of positivity for individual cell cycle regulators, ranging from 61.9 ± 2.0% up to 85.9 ± 1.2% for CDK4 and Cyclin D2 positive beta cells, respectively (Figure 1C). This discrepancy is probably due to differences in the half-life of the encoded proteins. For that reason, we could not accurately determine the overall IVT mRNA transfection efficiency in islet cells. Therefore, we used IVT mRNA encoding a highly stable green fluorescent protein (GFP) as a reporter marker. To assess IVT mRNA transfection efficiency in islet cells, we performed flow cytometry to measure both GFP and insulin positivity, the latter used as a beta-cell marker. After transfection with GFP IVT mRNA, the proportion of GFP positive beta cells reached 65.2 ± 2.0% at 24 hpt, further increasing up to 78.2 ± 3.7% at 36 hpt (Figures 1E and 1F).

### IVT mRNAs encoding cyclins D and CDKs induce cell cycle entry in rodent beta cells

Following the successful evaluation of IVT mRNA transfection and translation efficiency in islet cells, we investigated the effect of IVT mRNAs encoding cell cycle regulators on beta-cell proliferation. For this purpose, we used a cell proliferation assay based on the incorporation of thymidine analog 5-Ethynyl-2′-deoxyuridine (EdU) into newly synthesized DNA, combined with insulin immunostaining. This assay allowed us to detect beta cells that had entered the cell cycle and were progressing through the S phase, when DNA replication occurs. In addition, we adapted the EdU proliferation assay for flow cytometry-based analysis with the aim to more accurately quantify the proportion of EdU+ beta cells. We tested 12 pairs of cyclins D and CDKs IVT mRNAs, each consisting of CDK4 or CDK6 in combination with either WT or mutant forms of cyclin D1, D2, or D3. In addition, control samples of GFP IVT mRNA, transfection reagent-treated (Lipofectamine MessengerMAX), and untreated islet cells were included (Figure 2A).

**Figure 2.**
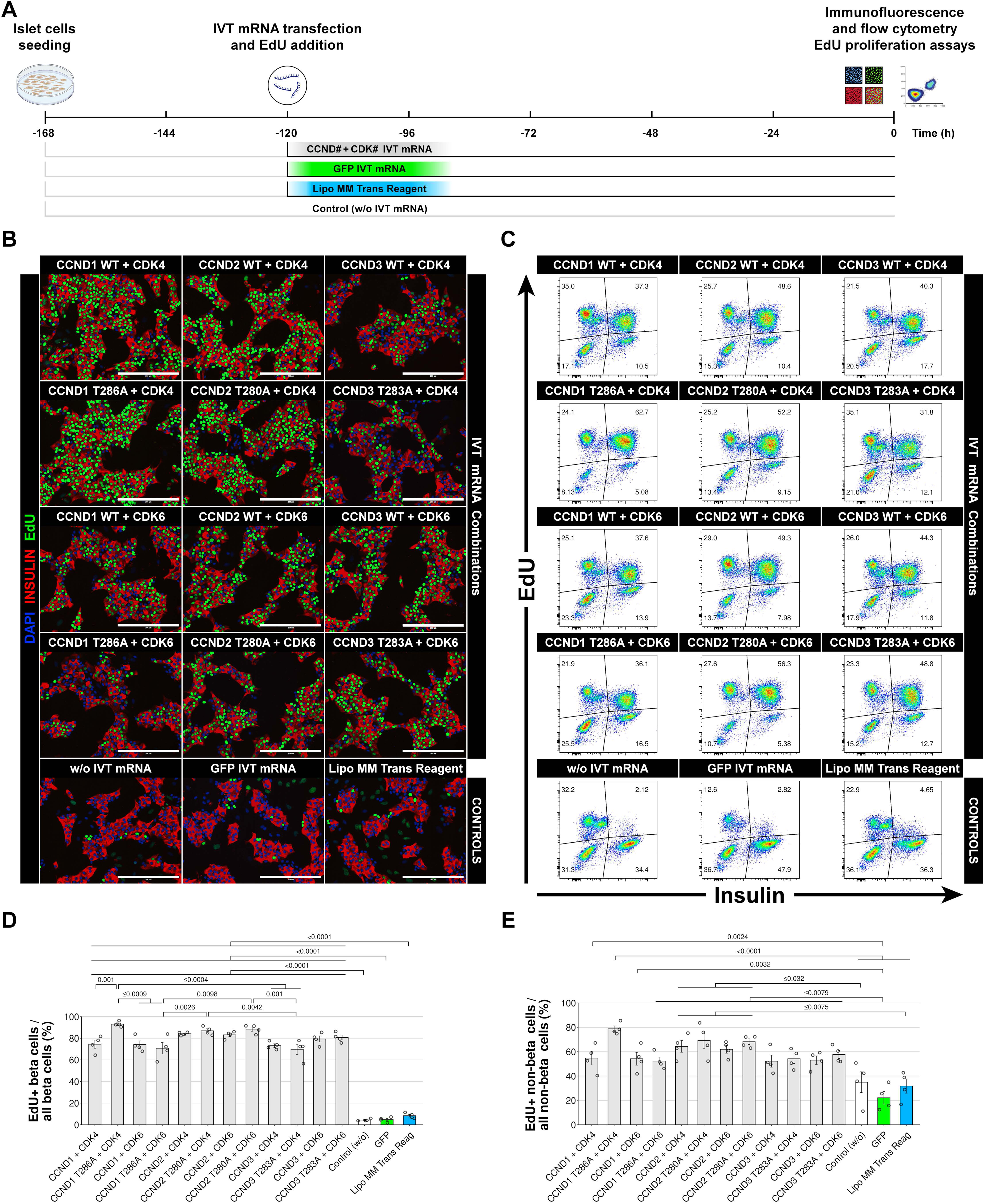
IVT mRNAs encoding cyclins D and CDKs activate cell cycle entry in rat beta cells. (A) Schematic showing the experimental design and timeline for testing IVT mRNA-induced beta-cell proliferation using immunofluorescence and flow cytometry EdU proliferation assays. (B) Representative images of co-stained insulin (red) and EdU (green) islet cells at 5 days after transfection with specific combinations of IVT mRNAs encoding cyclins D/CDKs. Nuclei are marked with DAPI (blue). GFP IVT mRNA treated, transfection reagent treated (Lipo MM Trans reagent), and untransfected islet cells were used as controls. Scale bars, 200 μm. (C) Representative flow cytometry plots and detection of EdU incorporation (Y-axis), and beta cells based on insulin intensity (X-axis), in IVT mRNA treated and control islet cell samples. (D) Quantification of EdU+ beta cells in IVT mRNA treated and control islet cell samples according to the flow cytometry detection of Edu+ incorporation. Data represent mean ± SD (n = 4 biologically independent samples per group). Statistical significance was determined using one-way ANOVA with Bonferroni’s post-test analysis. (E) Quantification of EdU+ non-beta-cells in IVT mRNA treated and control samples based on flow cytometry detection of Edu+ incorporation by non-beta-cells. Data represent mean ± SD (n = 4 biologically independent samples per group). Statistical significance was determined using one-way ANOVA with Bonferroni’s post-test analysis. P-values are shown above the graphs. Only significant values are shown.

We observed a significant increase in the number of EdU+ beta cells at 120 hpt across all samples treated with combinations of IVT mRNAs encoding cyclins and early CDKs using microscopic detection (Figure 2B) and flow cytometry analysis of EdU incorporation (Figure 2C). We detected the most significant increase in the proportion of EdU+ beta-cells (93.0 ± 1.2%) in samples treated with a mutant form of cyclin D1 (CCND1 T286A) together with CDK4 IVT mRNA (Figure 2D). This combination was also the only one in which we detected an advantageous effect of the mutant over the WT form of the cyclin D on the cell cycle activity of beta cells. CDK4 combined with WT CCND1 produced a significantly lower number of EdU+ beta cells (74.5 ± 3.8%). Surprisingly, we did not observe such a difference in samples treated with either mutant (CCND1 T286A) or WT CCND1 forms together with CDK6. These combinations activated comparable cell cycle entry rates (Figure 2D). The impact of a specific combination of IVT mRNAs on cell cycle activation was also evident in samples treated with Cyclin D3/CDKs IVT mRNA pairs. While islet cells treated with CDK4 combinations with either the WT CCND3 or mutant CCND3 T283A IVT mRNA exhibited one of the lowest rates of EdU+ beta cells, replacing CDK4 for CDK6 IVT mRNA in pairs containing CCND3 or CCND3 T283A IVT mRNA increased the proportion of EdU+ beta cells. Conversely, we observed the lowest variability in the proportion of EdU+ beta cells in samples treated with WT or mutant cyclin D2 forms in combination with either CDK. Moreover, these combinations of IVT mRNAs resulted in one of the highest rates of EdU+ beta cells out of all combinations tested. Notably, we detected a significantly lower rate of cell cycle activation in untreated control samples (4.3 ± 0.5% EdU+ beta cells).

In addition to cell cycle stimulation in beta cells, we observed a similar, though less pronounced, effect of IVT mRNAs on non-beta-cells (Figure 2E). Unsurprisingly, cell cycle activity in untreated control samples was substantially higher in the non-beta-cell fraction (34.9 ± 8.5% EdU+ non-beta-cells) than in beta cells (4.3 ± 0.5% EdU+ beta-cells).

Taken together, these data indicate that IVT mRNAs encoding cyclins D together with early CDKs can induce significant cell cycle activity in beta cells. In further experiments aiming to study cell cycle progression and the effects of IVT mRNA-induced beta-cell proliferation in more detail, we selected the most efficient combination of stimulatory IVT mRNAs, consisting of a mutant cyclin D1 form (CCND1 T283A) and CDK4, in Figures referred to as D1TAK4.

### Cell cycle dynamics of IVT mRNA-induced beta-cell proliferation

To gain further insight into the duration of IVT mRNA-induced beta-cell proliferation, we analyzed cell cycle activity using transient EdU labeling over a 120-hpt period. Specifically, we performed a pulse study by adding or removing the EdU analog from the media at various times after IVT mRNA transfection (Figure 3A). When we removed EdU at 12 h after IVT mRNA transfection, we observed non-significant DNA replication activity during the 0–12 hpt period (Figures 3B and 3C). As the study progressed, the number of EdU-labeled beta cells began to increase by 24 hpt, reaching 23.9 ± 1.3% EdU+ beta cells. The most notable increase occurred in samples co-incubated with EdU at 0–36 hpt, yielding 83.0 ± 1.1% EdU+ beta cells. Extending the incubation to 0–48 hpt and 0–72 hpt led to only slight additional increases in EdU labeling.

**Figure 3.**
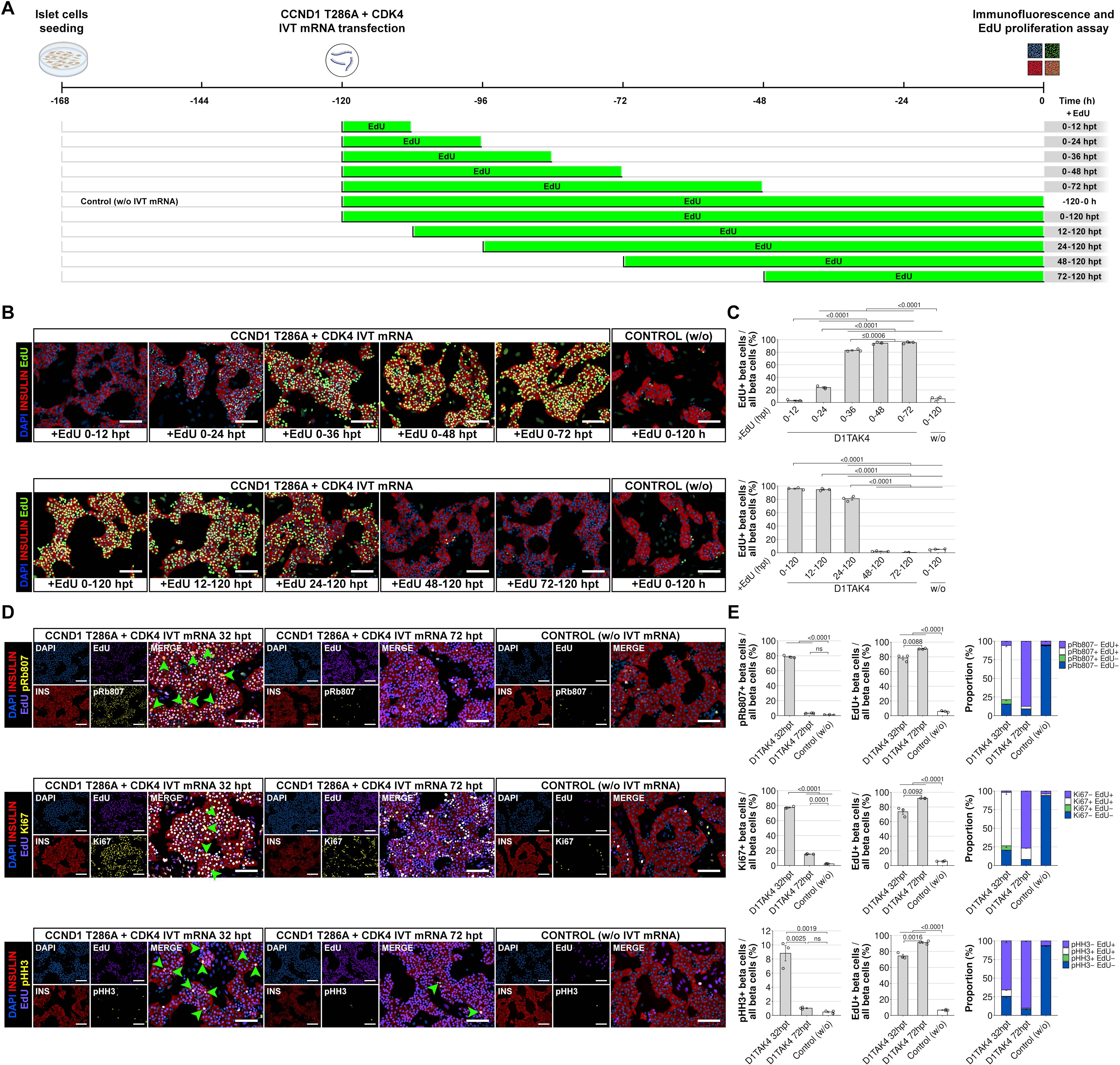
Analysis of cell cycle kinetics during the IVT mRNA induced beta-cell proliferation. (A) Schematic showing the experimental design of EdU removal/addition experiments after IVT mRNA transfection of rat islet cells. (B) Representative merged images of stained rat islet cells for insulin (red) and EdU (green) at 120 hpt after transfection with CCND1T286A and CDK4 IVT mRNA, followed by EdU coincubation for specified period (indicated at the bottom of each image). Nuclei are marked with DAPI (blue). (C) Corresponding high-content imaging-based quantification of EdU+ beta cells, related to EdU coincubation time, following CCND1T286A and CDK4 IVT mRNAs transfection (abbreviated as D1TAK4). Data represent mean ± SD (n = 3 biologically independent samples per group). Statistical significance was determined using one-way ANOVA with Bonferroni’s post-test analysis. P-values are shown above the graphs. Only significant values are shown. (D) Representative images of rat islet cells stained for insulin (red), EdU (purple), and cell cycle markers (pRb807; Ki67; pHH3) (yellow) at 32 and 72 hpt and treated with CCND1T286A and CDK4 IVT mRNA. Nuclei are marked with DAPI (blue). Green arrowheads indicate beta-cell mitotic nuclei. Untransfected islet cells were used as control samples. Scale bars, 100 μm. (E) High-content imaging-based quantification of pRb807+; Ki67+; pHH3+; and EdU+ beta-cells (left and middle), and the corresponding proportions of beta-cell subpopulations based on specific cell cycle marker/EdU positivity (right), in CCND1T286A and CDK4 IVT mRNA treated samples (abbreviated as D1TAK4), and control untreated samples. Data represent mean ± SD (n = 3 biologically independent samples per group). Statistical significance was determined using one-way ANOVA with Bonferroni’s post-test analysis. Colors on bar charts correspond to the specific nuclei colors shown on merged images. P-values are shown above the graphs. Only significant values are shown.

Next, we confirmed and extended these results by EdU addition experiments, where we added an EdU analog into the media at different time points after IVT mRNA transfection (Figure 3A). Unsurprisingly, EdU labeling was highest when we applied EdU during the early post-transfection period at 0 and 12 hpt, with 96.0 ± 0.9% and 94.6 ± 1.0% EdU+ beta cells, respectively (Figures 3B and 3C). EdU addition at a later time point (24 hpt) caused a slight decline in EdU positivity, with 81.0 ± 2.2% EdU+ beta cells. Nevertheless, we detected the most significant decrease in EdU labeling when applying EdU from 48 hpt onward, with only 2.2 ± 0.4% EdU+ beta cells. This decline was even more evident when we added EdU into the media at 72 hpt.

To better assess the progression of beta cells through the entire cell cycle, we performed immunostaining for cell cycle-specific markers at various time points after mRNA transfection. We analyzed islet cells for the phosphorylated form of the retinoblastoma protein (pRb807/811; marking cycling cells from an early G1 phase until mitosis), the widely used cell cycle marker Ki67 (expressed throughout the whole cell cycle, except early G1), and the phosphorylated form of histone H3 (pHH3; marking cells in mitosis).

Immunostaining for pRb807 (Figures 3D, 3E, and S1A) revealed that some beta cells enter the cell cycle as early as 4 h following IVT mRNA transfection. Nevertheless, most beta cells entered the cell cycle within the first 12 hpt, with the maximal proportion of pRb807-positive beta cells detected at 32 hpt (78.7 ± 1.0% pRb807+ beta cells). We confirmed these results by additionally detecting the cell cycle marker Ki67 (Figures 3D, 3E, and S1B), whose expression was slightly delayed compared to pRb807. First, we observed an increased Ki67+ beta-cell numbers at 12 hpt, with a significant further increase up to 32 hpt (77.6 ± 0.6% Ki67+ beta cells). During the following period, beta-cell positivity for both these cell cycle markers declined at 72 hpt. Finally, at 96 hpt, the number of pRb807+ and Ki67+ beta cells declined to levels comparable with untreated control samples.

Additional analysis of the mitotic marker pHH3 (Figures 3D, 3E, and S1C) revealed increased pHH3 positivity as early as 24 hpt. We observed the highest percentage of pHH3+ beta cells at 32 hpt (8.8 ± 1.1% pHH3+ beta cells). During that period, we also observed a high occurrence of morphological changes in beta-cell nuclei resembling cytokinesis (green arrows in Figure 3D), confirming ongoing cell division. Later, the number of pHH3+ beta cells gradually declined to 0.4 ± 0.1% at 96 hpt. These numbers reached levels comparable to untreated controls, confirming the results from pRb807 and Ki67 immunostaining and indicating completed cell division and a return to quiescence.

Together, these results suggest that after IVT mRNA treatment, some beta cells enter the G1 phase as early as 4 hpt, while most follow within 12 hpt. DNA synthesis (S phase) begins in active cells during 12– 24 hpt, peaking for the majority between 24–36 hpt. Rapidly proliferating beta cells initiate mitosis around 24 hpt, though peak mitotic activity occurs at ∼32 hpt. Finally, a significant decline in cell cycle activity at 72 hpt is accompanied by the subsequent cessation of IVT mRNA–induced proliferation.

### IVT mRNA-induced cell cycle entry results in beta-cell division and increases beta-cell mass

Although cell cycle markers provide important information about ongoing proliferative activity, they do not offer direct evidence of a completed cell cycle and division. To investigate whether the stimulatory IVT mRNA can not only initiate a cell cycle but also lead to successful cell division, we employed two independent assays that provide direct evidence of cell division.

The first method combines FACS-based beta-cell sorting with high-content imaging to measure changes in total beta-cell numbers. At first, we sorted a defined number of beta cells (30,000) using the zinc ion-specific fluorophore FluoZin-3, which specifically labels pancreatic beta cells. Next, we treated purified beta cells with the stimulatory IVT mRNAs (CCND1 T283A + CDK4) and cultured them for four days. Finally, we performed insulin and EdU co-staining to quantify changes in beta-cell numbers and EdU incorporation (Figures 4A and 4B). FluoZin-3-based cell sorting allowed us to obtain a highly pure population of beta cells with > 99% purity, as determined by the proportion of insulin+/DAPI+ cells. Using high-content imaging analysis, we detected an almost 2-fold increase in the total number of beta cells (48,407 ± 2,912 versus 24,654 ± 783), four days after stimulatory IVT mRNA treatment (Figure 4B). The increased beta-cell numbers were accompanied by a high proportion of EdU+ beta cells in IVT mRNA-treated samples (Figure 4B).

**Figure 4.**
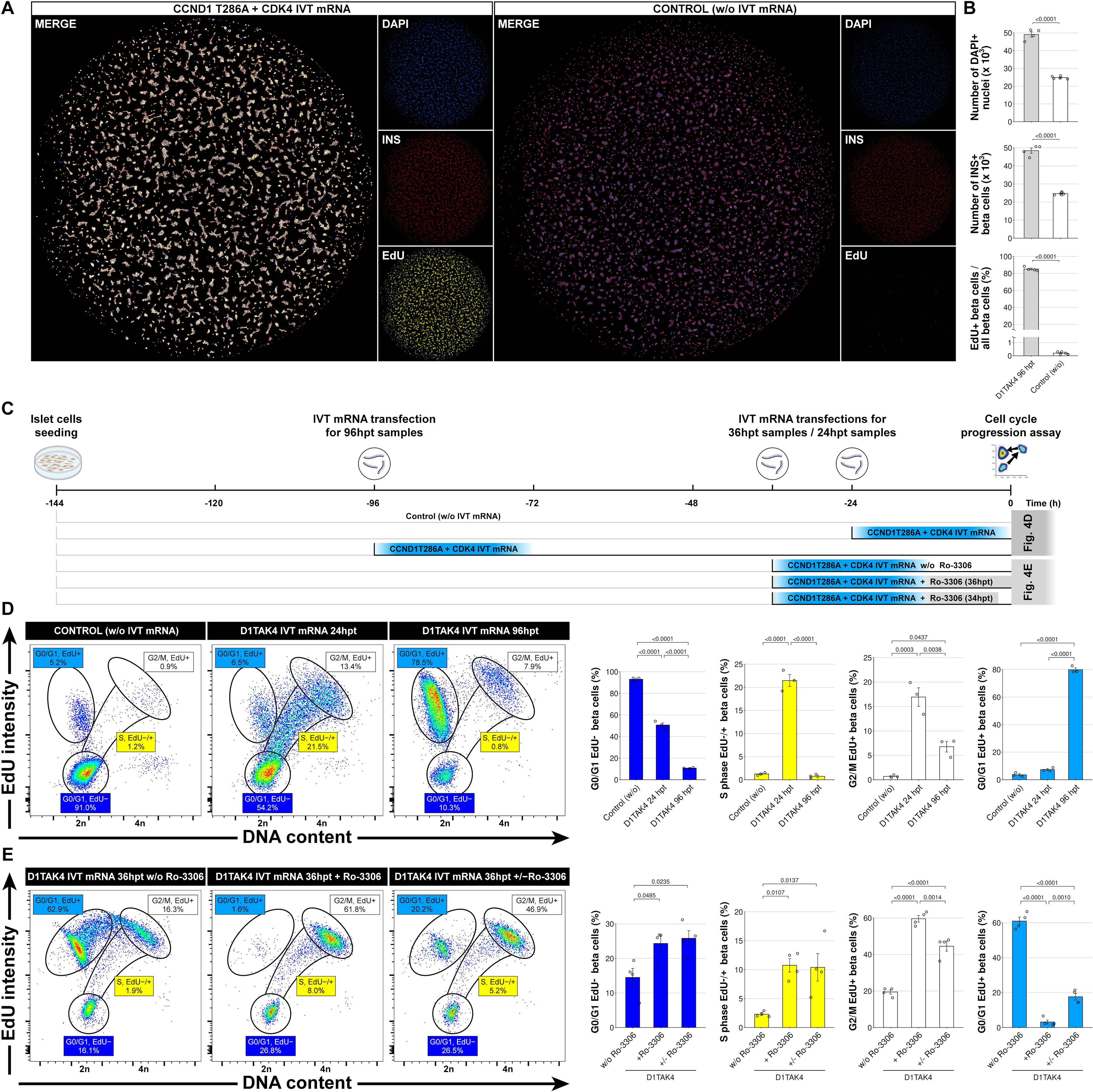
Stimulatory IVT mRNAs induce cell cycle entry, cell division, and increased beta-cell mass. (A) Representative tiled images covering complete 96-well plates seeded with FACS-sorted beta cells (30,000/well). Samples transfected with IVT mRNAs or untreated controls were stained for cell nuclei (DAPI), insulin (INS), and EdU. (B) High-content imaging-based quantification of the total cell number (DAPI), beta-cell number (INS), and proportion of EdU+ beta cells in IVT mRNA-treated samples (abbreviated as D1TAK4) as well as untreated controls. Data represent mean ± SD (n = 4 biologically independent samples per group). Statistical significance was determined using two-sided Student’s t-test. P-values are shown above the graphs. (C) Schematic showing the experimental design of the cell cycle progression assay. Islet cells were cultured for 6 days and treated at different time points with stimulatory IVT mRNAs alone or together with a CDK1 inhibitor Ro-3306. At the end, cells were harvested and processed for a flow cytometry-based cell cycle assay. (D) Representative flow cytometry plots and quantification of beta-cell-specific cell cycle subpopulations in untreated control and IVT mRNA treated (24 and 96 hpt) samples, using the cell cycle progression assay. Circles and S-phase corridor on the plots indicate specific cell cycle phases/gated populations used for analysis. Data represent mean ± SD (n = 3 biologically independent samples per group). Statistical significance was determined using one-way ANOVA with Bonferroni’s post-test analysis. P-values are shown above the graphs. Only significant values are shown. (E) Verification of cell cycle assay and quantification of beta-cell-specific cell cycle subpopulations, using CDK1 inhibitor Ro-3306, in IVT mRNA treated (36 hpt) islet cells samples. Included is a sample where Ro-3306 was washed out during the last 2 h of incubation (abbreviated as D1TAK4 IVT mRNA 36 hpt +/− Ro3306). Data represent mean ± SD (n = 4 biologically independent samples per group). Statistical significance was determined using one-way ANOVA with Bonferroni’s post-test analysis. P-values are shown above the graphs. Only significant values are shown.

Next, we used a flow cytometry-based cell cycle progression assay to confirm that stimulatory IVT mRNAs ultimately induce beta-cell division. The cell cycle progression assay simultaneously detects DNA content and EdU incorporation (DNA synthesis). Simultaneous use of these two parameters allows quantification of cell populations at different cell cycle phases, including cells that have completed mitosis.^32^

Using this assay, we confirmed the results from the IF experiments. An increased number of beta cells entering and progressing through the cell cycle was accompanied by a decline in the number of quiescent and early G1-phase (G0/G1 EdU-) beta cells (Figure 4D). Following stimulatory IVT mRNA transfection (at 24 hpt), we detected a significant increase in the proportion of S phase, and G2/M beta cells, together with a decline in the number of G0/G1 EdU-beta-cells. However, at that time point, we did not observe a notable increase in the number of divided beta cells (G0/G1 EdU+). Nevertheless, at the next time point (36 hpt), we detected substantial progression throughout the cell cycle as evidenced by an increased number of divided beta cells (61.0 ± 2.3% G0/G1 EdU+) and an additional decline in the number of G0/G1, EdU-beta cells. Finally, at 96 hpt, the total proportion of divided beta cells increased up to 80.0 ± 1.5%, while only 10.8 ± 0.3% beta cells remained in the quiescent G0/G1 EdU-fraction (Figure 4D). Furthermore, cell cycle attenuation at that time point was confirmed by the decreased number of beta cells in the S- and G2/M-phases.

In addition, we confirmed that the transition between G2/M and G0/G1 EdU+ populations captures a real cell division. Thus, we used a CDK1-specific inhibitor (Ro-3306), widely used in cell cycle/mitotic studies.^33^ Ro-3306 inhibits CDK1, which causes reversible G2/M cell cycle arrest. In samples treated with a combination of stimulatory IVT mRNAs and Ro-3306, we detected a significant increase in the number of undivided G2/M, and a simultaneous decline in the divided G0/G1 EdU+ beta-cell populations at 36 hpt (Figure 4E). We further demonstrated that beta cells can resume mitosis and complete cell division after the release of the mitotic block. Indeed, at 2 h after Ro-3306 washout (Figure 4E, denoted as D1TAK4 IVT mRNA 36 hpt +/−Ro-3306), we observed a significant increase in the number of divided beta cells, accompanied by a decrease in the undivided beta-cell subpopulations.

In sum, the results from the cell cycle progression assay indicate that stimulatory IVT mRNAs not only promote cell cycle entry but also facilitate successful cell division in rodent beta cells. FACS-based high-content imaging assay confirmed these results, providing further evidence that IVT mRNA-induced proliferation is associated with an almost twofold increase in the total beta-cell numbers.

### Gene expression profile during IVT mRNA-induced islet cell proliferation

To characterize changes during IVT mRNA-induced proliferation more precisely, we analyzed islet cell gene expression at defined times after transfection with stimulatory IVT mRNAs (Figure 5A). Based on previous cell cycle progression experiments, we chose 24 hpt as the initial time point. At this stage, most beta cells had entered the cell cycle and were present in all phases, but had not yet undergone extensive division. We also included samples from 48 hpt, when beta-cell proliferation was still ongoing, even though most beta cells had completed cell division. Finally, we analyzed islet cells at 120 hpt, when IVT mRNA-induced proliferation should have finished, and most beta cells were expected to be quiescent.

**Figure 5.**
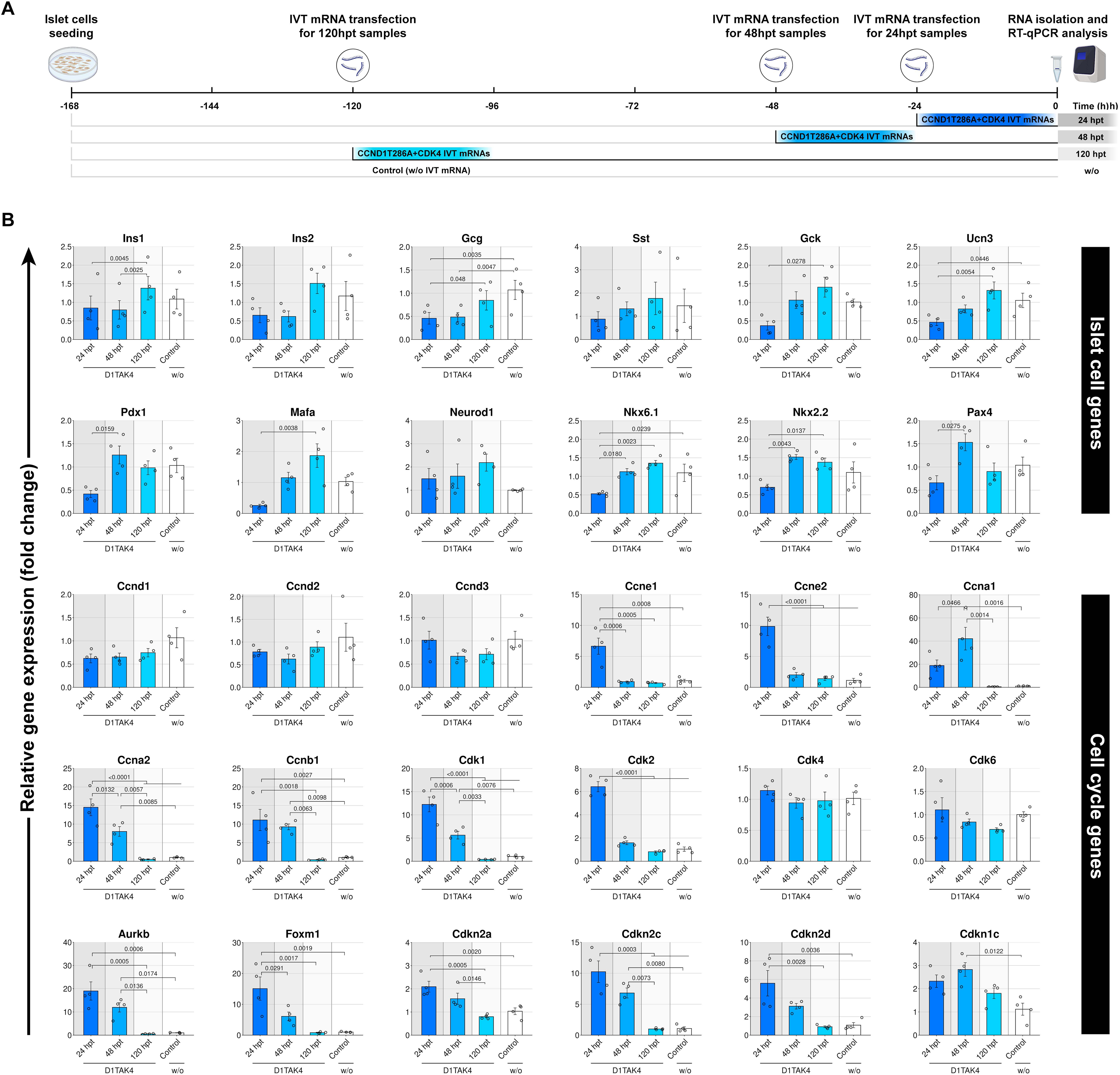
Gene expression changes during IVT mRNA-induced islet cell proliferation. (A) Schematic of the experiment for testing the effect of IVT mRNA-induced proliferation on islet cell gene expression. Islet cells were transfected with stimulatory IVT mRNAs at different time points (day 2, 5, 6) during a 7-day culture period. Samples were collected on day 7, total RNA purified, and analyzed by RT-qPCR. (B) RT-qPCR analysis of genes involved in cell cycle regulation and islet cells/beta-cell function at 24; 48; 120; hpt. Values indicate fold change relative to the expression levels of untreated control samples (normalized to *Hprt* and *Actb* expression). Data represent mean values ± standard error of the mean (SEM), and scatterplots show all individual data points. Statistical significance was determined using two-way ANOVA followed by Bonferroni’s post-test analysis; n = 4 biologically independent samples per group. P-values are shown above the graphs. Only significant values are shown.

RT-qPCR analysis showed a significant increase in endogenous expression of cell cycle-related genes at 24 hpt (Figure 5B). These included genes encoding cyclins (E1, E2, A1, A2, B1), CDKs (*Cdk1* and *Cdk2*), and other regulators such as *FoxM1* and *Aurkb*, participating in intermediate and late cell cycle stages. Additionally, cell cycle inhibitors such as *Cdkn2a/c/d* and *Cdkn1c* were also upregulated in IVT mRNA-treated samples. This result is in agreement with previous studies of induced beta-cell proliferation, showing an induction of some cell cycle inhibitors upon ectopic overexpression of cell cycle activators.^34^ As expected, the only cell cycle-related genes not upregulated after stimulatory mRNA treatment were those involved in the early G1 phase, such as *Ccnd1*, *Ccnd2*, *Ccnd3*, *Cdk4*, and *Cdk6*. During the following period, expression of most cell cycle genes declined significantly, except for *Ccnb1* and *Aurkb*, whose expression remained unchanged between 24 and 48 hpt, and *Ccna1*, which even increased further.

Changes in the expression of islet cell-related genes and beta-cell transcription factors (TFs) were less pronounced. At 24 hpt, we observed a slight, statistically non-significant downregulation of genes encoding pancreatic hormones (*Ins1*, *Ins2*, *Gcg*, *Sst*) and some important beta-cell-specific TFs (*Mafa*, *Nkx6.1* and *Pdx1*). Nevertheless, their expression reversed and returned to normal levels in the following period. This result is in agreement with a previous study reporting that *Pdx1* expression is highly variable throughout the beta-cell cycle, with significant downregulation during the S phase.^35^ Notably, the expression of the beta-cell maturity marker *Ucn3* also declined at 24 hpt, indicating potential beta-cell dedifferentiation during the early phases of IVT mRNA-induced proliferation.

Consistent with previous analyses, the expression of cell cycle and islet/beta-cell-related genes normalized within 120 hpt, marking the end of IVT mRNA-induced proliferation and a return to quiescence.

### Beta-cell-specific gene expression changes during IVT mRNA-induced proliferation

Gene expression analysis revealed highly dynamic changes in the expression of cell cycle-related genes during IVT mRNA-induced islet cell proliferation. However, we obtained these results from an unpurified islet cell population containing various types of endocrine and non-endocrine cells. To gain a more detailed insight into beta-cell-specific gene expression changes, we took advantage of Fluozin-3-based FACS sorting and performed a bulk RNA sequencing (RNA-Seq) of purified beta cells.

RNA-Seq was performed on purified beta-cells at 1 or 5 days after transfection with stimulatory or GFP IVT mRNAs (Figure 6A). As determined in previous cell cycle progression experiments, samples obtained at 1 day after stimulatory mRNA transfection contained mainly beta cells in all cell cycle phases, while cells obtained at 5 days after IVT mRNA transfection included quiescent beta cells with an accomplished cell cycle.

**Figure 6.**
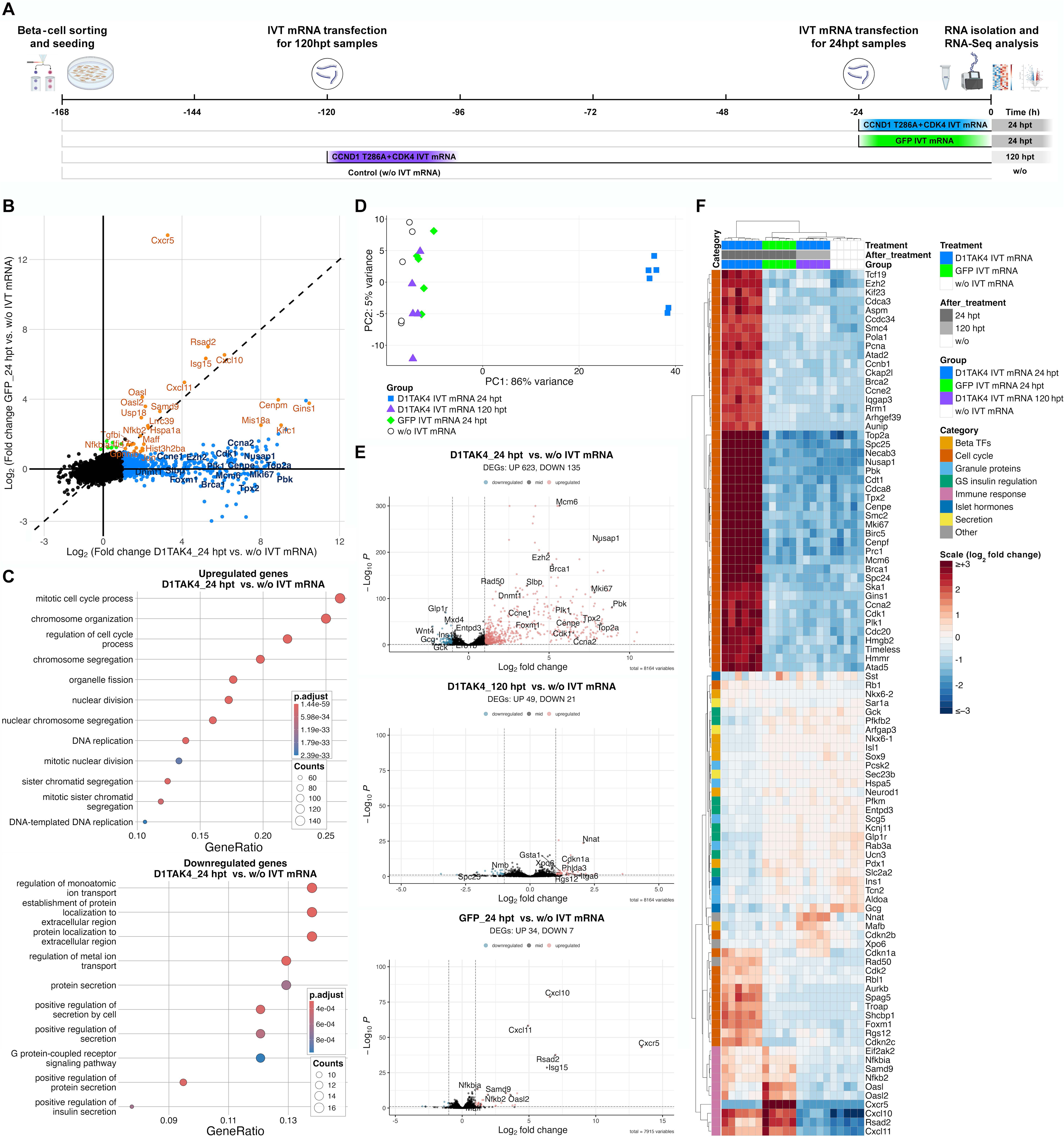
RNA-seq analysis of purified beta-cells during IVT mRNA induced proliferation reveals upregulation of cell cycle related genes. (A) Schematic showing the experimental design of RNA-seq analysis and division into sample subgroups based on IVT mRNA transfection. (B) Scatterplot showing differentially expressed genes (DEGs) in purified beta-cells at 24 hpt with either GFP IVT mRNA (y-axis) or CD1TA/CDK4 IVT mRNAs (x-axis) treatment, compared to untreated control samples (p adjusted < 0.05). Each dot represents a unique gene; green circles denote log2 (fold change) >1, upregulated genes in GFP IVT mRNA treated beta-cells; blue circles denote log2 (fold change) >1, upregulated in CD1TA/CDK4 IVT mRNAs treated beta-cells, orange circles denote log2 (fold change) >1, upregulated in both CD1TA/CDK4 and GFP IVT mRNAs treated beta cells, and black indicates unchanged genes. Selected statistically significant upregulated and downregulated genes, as determined by a two-sided Chi-Square test, are indicated. (C) Over-representation analysis (ORA) showing the most upregulated and the most downregulated pathways in CD1TA/CDK4 IVT mRNAs treated beta cells (24 hpt) categorized based on gene ontology analysis. Gene sets with FDR < 0.05 and P-value < 0.05 were considered significant. (D) Principal component analysis of RNA-Seq data showing distinct molecular profiles during and after IVT mRNA induced beta-cell proliferation. Sample groups treated with CD1TA/CDK4 IVT mRNAs (24 hpt and 120 hpt), GFP IVT mRNA (24 hpt), and untreated control sample of purified beta cells, indicated by different colors. n = 5 samples for each group, 50 000 beta-cells per sample. (E) Volcano plots showing differentially expressed genes (DEGs) (P adjusted < 0.05 (displayed as - log10), and log2 fold change <−1 or log2 fold change >1) in sorted beta-cells at 24 hpt with CD1TA/CDK4 IVT mRNA, at 24 hpt with GFP IVT mRNA, and 120 hpt with CD1TA/CDK4 IVT mRNAs, all compared to untreated controls. Downregulated genes are shown in blue (log2 (fold change) <−1), upregulated genes in red (log2 (fold change) >1), and unchanged genes in black. (F) Heatmap and hierarchical clustering analysis showing key genes associated with IVT mRNA induced beta-cell proliferation. The values in the heatmap are log2-scaled RNA-seq read counts. The color scale indicates the gene expression level: red denotes log2 (fold change) > 3, blue log2 (fold change) < −3. n = 5 experiments per condition.

The most significantly induced genes in replicating beta-cells (24 hpt) included cell cycle-related genes, particularly those involved in regulation of cell cycle progression and mitotic processes, chromosome organization and segregation, organelle fission, nuclear division, DNA replication and mitotic nuclear division, and sister chromatid segregation, as revealed by gene ontology (GO) analysis (Figure 6C). Significant upregulation of S- and G2/M-phases specific cyclins and CDKs such as *Ccne2*, *Ccna2*, *Ccnb1*, *Cdk1*, *Cdk2* as well as *Foxm1* and *Aurkb* confirmed RT-qPCR results (Figures 6E and 6F). In addition, among the most significantly overexpressed genes were those previously identified during the analysis of beta-cell proliferation in vivo,^36,37^ including *Top2a*, *Plk1*, *Nusap1*, *Pbk*, *Cdk1*, *Cenpe*, *Cenpf*, *Tpx2*, *Hmmr*, *Cdca3*, *Cdca8*, *Birc5*, *Ska1*, and *Aspm*. On the other hand, genes downregulated during IVT mRNA-induced proliferation were associated with processes important for beta-cell function, mainly protein secretion and regulation of signal release, such as *Rab3a*, *Gck*, *Slc2a2*, *and Glp1r* (Figures 6C and 6F). RNA-Seq analysis also confirmed downregulation of beta-cell-specific TFs Pdx1 and Nkx6.1, maturation markers *Ucn3*, *Entpd3*,^38^ and *Ins1*, altogether suggesting partial dedifferentiation during mRNA-induced beta-cell replication. Interestingly, *Nkx6.1* TF downregulation was accompanied by upregulation of its paralog *Nkx6.2*, potentially acting as a compensatory mechanism.^39^

To gain further insights into the potential side effects of the IVT mRNA treatment itself, we analyzed samples of purified beta cells at 1 day after GFP IVT mRNA transfection. Both GFP and stimulatory IVT mRNA transfections resulted in upregulation of innate immune response genes such as C–X–C motif chemokine 10 (*Cxcl10*; also known as IFNγ-induced protein 10), C–X–C motif chemokine 11 (*Cxcl11*), and *Rsad2*, along with *Isg15*, both interferon-stimulated genes (Figure 6F). Of particular interest for the current study was finding that the upregulation of some innate immune genes in beta cells was related to the specific IVT mRNA used. We observed higher *Cxcr5*, *Oasl*, and *Oasl2* expression in samples treated with GFP IVT mRNA than in samples treated with stimulatory IVT mRNAs (Figure 6D and 6F).

### Stimulation of human beta-cell proliferation by IVT mRNAs encoding cyclin D1, CDK4, and MYC

Finally, we investigated whether the stimulatory IVT mRNAs can also induce the proliferation of human-derived beta cells. To that end, we tested a combination of IVT mRNAs encoding cell cycle regulators on EndoC-BH5 cells. These cells are derived from human fetal beta cells and currently represent the only model of human beta cells available in sufficient quantity and standardized quality. These cells are quiescent and have an insulin content and insulin secretory capacity comparable to those of native human beta cells.^40^

Our preliminary results showed that human EndoC-BH5 cells can enter and progress through the cell cycle; however, they cannot complete cell division when treated with a simple combination of IVT mRNAs encoding human CCND1 T286A and CDK4 (Figure S2). To promote cell division in EndoC-BH5 cells, we applied an additional IVT mRNA encoding the MYC gene, enabling successful completion of the cell cycle and mitotic division.

To gain further insight into the IVT mRNA-induced EndoC-BH5 proliferation, we performed IF staining of cell cycle markers (pRb807, Ki67, CCNA2, and pHH3), along with EdU detection over a 96-hpt period (Figure 7A). This showed that a combination of CCND1 T286A, CDK4, and MYC IVT mRNAs induced cell cycle entry in most EndoC-BH5 cells within a 60-hpt period. Further, we detected a significant increase in the numbers of pRb807+, Ki67+, CCNA2+, and pHH3+ EndoC-BH5 cells compared with untreated controls at 60 hpt (Figure 7B). A similar increase in EdU positivity accompanied high cell cycle activity. In the next 36 h (at 96 hpt), we observed a substantial decline in positivity for most cell cycle markers, including pRb807+, CCNA2+, and pHH3+. These numbers reached levels comparable with untreated controls, except for Ki67+ cells, which remained elevated even at 96 hpt.

**Figure 7.**
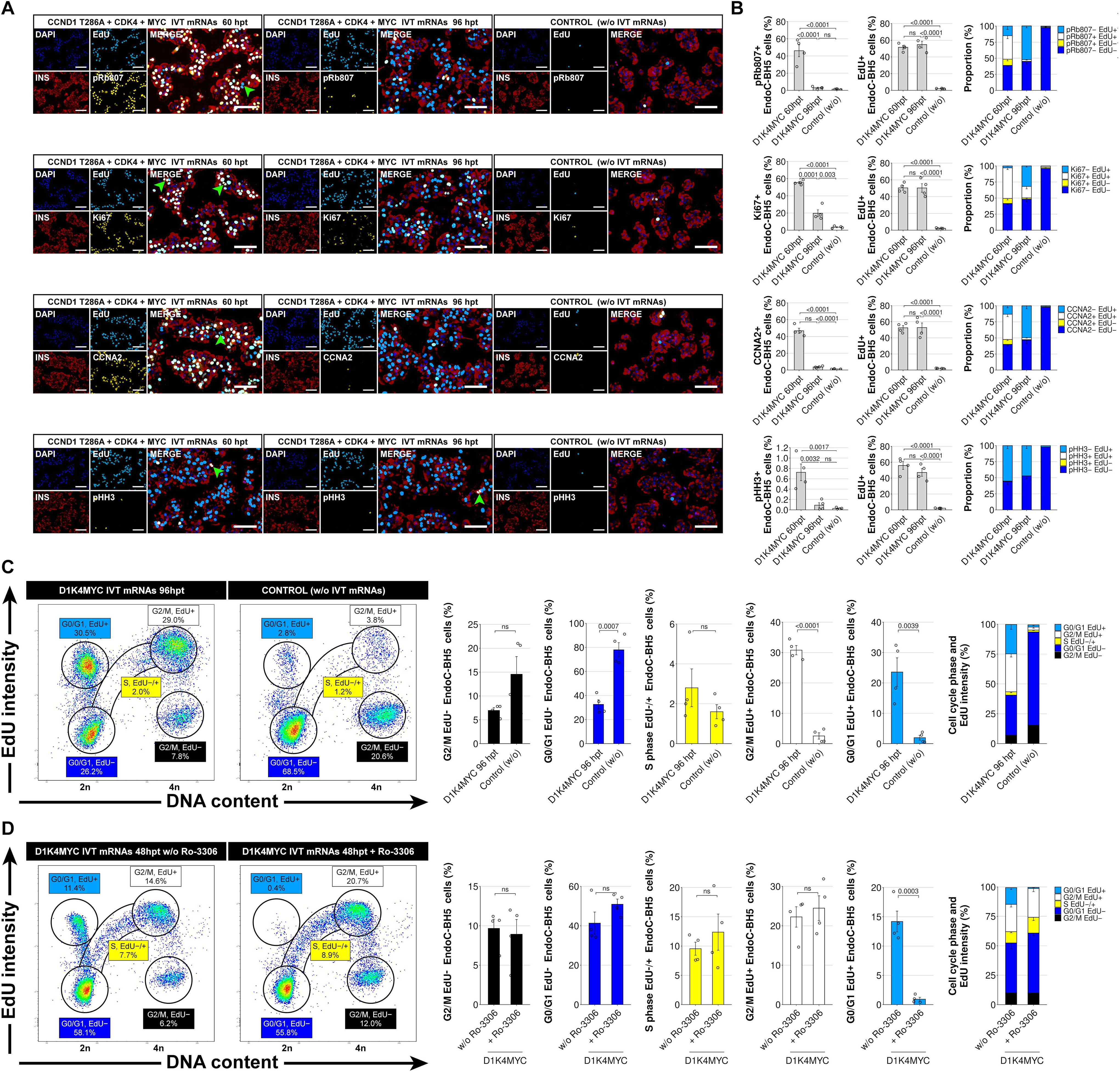
IVT mRNAs encoding cyclin D, CDK4 and MYC activates cell cycle entry and mitosis in human fetal beta-cell derived EndoC-BH5 cells. (A) Representative images of insulin (red), EdU (cyan), and cell cycle marker (pRb807; Ki67; CCNA2; pHH3) (yellow) staining, of EndoC-BH5 cells at 60 and 96 hpt treated with a combination of CCND1T286A, CDK4, and MYC (D1K4MYC) IVT mRNAs. Nuclei are marked with DAPI (blue). Green arrowheads indicate mitotic nuclei. Untransfected EndoC-BH5 cells were used as controls. Scale bars, 100 μm. (B) Quantification of pRb807+; Ki67+; CCNA2+; pHH3+; and EdU+ EndoC-BH5 cells and the proportions of respective subpopulations based on specific cell cycle marker/EdU positivity, in IVT mRNA treated and control untreated samples according to high-content imaging-based quantification. Data represent mean ± SD (n = 4 biologically independent samples per group). Statistical significance was determined using one-way ANOVA with Bonferroni’s post-test analysis. P-values are shown above the graphs. Colors on bar charts correspond to specific nuclei colors on merged images. (C) Representative flow cytometry plots and quantification of cell cycle subpopulations in IVT mRNA treated (96 hpt) and control untreated EndoC-BH5 cell samples, based on the cell cycle progression assay. Circles and S-phase corridor indicate the gated populations used for analysis. Data represent mean ± SD (n = 4 biologically independent samples per group). Statistical significance was determined using two-sided Student’s t-test. P-values are shown above the graphs. (D) Verification of the cell cycle progression assay and quantification of cell cycle subpopulations using CDK1 inhibitor Ro-3306, in IVT mRNA treated (48 hpt) EndoC-BH5 cell samples. Data represent mean ± SD (n = 4 biologically independent samples per group). Statistical significance was determined using two-sided Student’s t-test. P-values are shown above the graphs.

To confirm and extend the results of the IF/EdU analysis, we performed the cell cycle progression assay on EndoC-BH5 cells treated with a combination of stimulatory IVT mRNAs (Figures 7C and 7D). Despite the delayed cell cycle progression of human EndoC-BH5 cells compared to rodent beta cells, 14.2 ± 1.8% of EndoC-BH5 cells had already undergone mitosis and divided at 48 hpt (Figure 7D). At this time point, there was ongoing proliferative activity, evidenced by the increased proportion of cells in S and G2/M-phases. At 96 hpt, we observed an additional increase in the number of divided cells (23.6 ± 4.7% G0/G1 EdU+) (Figure 7C) and an attenuation of cell cycle activity with a notable decline in the number of cells remaining in the S phase. Surprisingly, the number of EndoC-BH5 cells at G2/M further increased up to 30.8 ± 1.5% G2/M EdU+ cells at 96 hpt. This result is consistent with persistent Ki67 positivity, as detected by IF analysis at 96 hpt.

Interestingly, in all samples examined, we detected a subpopulation of EdU-negative EndoC-BH5 cells in the G2/M phase (G2/M EdU−). Since we applied the EdU analog throughout the entire test period, this G2/M cell population had to be present in the cell culture already at the beginning. Moreover, since this subpopulation was also found in untreated control samples, it is presumably an artifact generated during the EndoC-BH5 cell preparation, rather than a result of IVT mRNA treatment.

Finally, we confirmed that the transition between G2/M EdU+ and G0/G1 EdU+ populations effectively captures the cell division of EndoC-BH5 cells (Figure 7D). Addition of the CDK1 inhibitor Ro-3306 caused a significant blockade of mitotic activity, as shown by a decreased number of divided G0/G1 EdU+ EndoC-BH5 cells at 48 hpt.

These results collectively imply that IVT mRNAs encoding cell cycle regulators can induce proliferation of human-derived beta cells. Accordingly, we propose that most EndoC-BH5 cells divide during the first 60 h after IVT mRNA transfection. We draw this conclusion from a stable proportion of EdU-positive cells observed between 60 hpt and 96 hpt, alongside a simultaneous decline in the numbers of pRb807+, Ki67+, CCNA2+, and pHH3+ cells during the same period. While many EndoC-BH5 cells completed the cell cycle and divided, a similar proportion remained undivided in the G2/M EdU+ state. We suppose that, in addition to other possible factors, the reluctance of some EndoC-BH5 cells to complete the cell cycle may be due to insufficient induction of key cell cycle regulators or induced cell cycle arrest.

## DISCUSSION

Our study introduces a novel strategy for inducing pancreatic beta-cell proliferation using IVT mRNA encoding specific cell-cycle regulators. This approach could activate cell cycle entry and cell division in both rodent- and human-derived beta cells. To the best of our knowledge, these results represent the first evidence of IVT mRNÁs applicability for the stimulation of beta-cell proliferation. Direct and indirect observations support the accomplishment of cell cycle and successful beta-cell division. These include the detection of beta-cell mitotic nuclei and the mitotic marker pHH3 during periods of intense cell cycle activity, an almost twofold increase in beta-cell number in purified beta-cell cultures, and accomplished beta-cell division, as confirmed by the cell cycle progression assay.

We could obtain these results mainly due to the unique properties of IVT mRNA, including significant yet transient overexpression of the encoded protein and minimal immunogenicity.^14,21^ One of the major advantages of the IVT mRNA lies in its temporal expression profile, which is intrinsic to the nature of mRNA. The precise control of expression kinetics is particularly critical for cellular processes, such as the immune response and cell cycle regulation, which rely on rapid and dynamic changes in the expression levels of regulatory proteins.^10,12,41^ Additionally, IVT mRNA provides a significant overexpression of a specific protein within a relatively short timeframe. For instance, the dose-dependent effect of cyclinD1/CDK6 expression on human beta-cell proliferation was previously reported by Takane et al.^11^ The current study is an additional confirmation of our own and others’ previous findings, highlighting the potential of IVT mRNA technology for cell fate manipulation, gene expression regulation, and gene editing.^25,42–44^

Our study also showed that the extent and progression of beta-cell proliferation depend not only on the presence of cell cycle regulators but also on their specific combination. This was true for both rodent and human beta cells. In rodent beta cells, altering just one partner in cyclins D/CDK pairs led to a >30% difference in the proportion of EdU+ cells. In human EndoC-BH5 cells, the effect of particular cell cycle regulators on the overall cell cycle output was even more striking. While CCND1 T286A and CDK4 IVT mRNAs alone could induce entry and progression through the cell cycle, they could not promote cell division. To achieve this, we had to include an additional IVT mRNA encoding MYC. This result aligns with prior studies showing that co-application of late-stage cell cycle regulators, such as CDK1 and cyclin E, or cyclin B, in addition to early cell cycle regulators, significantly increases overall proliferation of human beta-cells^45^ and cardiomyocytes.^46^

The essential role of c-MYC in beta-cell proliferation is well established. For instance, Karslioglu et al. demonstrated that modest c-MYC overexpression can overcome the intrinsic proliferative silence of human beta cells.^47^ Beyond its role in proliferation, c-MYC also regulates other cellular processes critical for proper cell cycle initiation and progression, including metabolism, DNA repair, epigenetic modifications, dedifferentiation, and apoptosis. Its upregulation is known to support compensatory beta-cell expansion in response to an increased metabolic demand.^48^ In addition, Puri et al. reported that the positive effect of c-MYC on beta-cell proliferation is associated with a partial shift toward an immature phenotype, reminiscent of the beta-cell state soon after birth, which has enhanced proliferative potential.^49^ Conversely, silencing endogenous c-MYC reduced beta-cell proliferative capacity. These observations reinforce the potential of IVT mRNA for transient c-MYC expression: its short-lived nature enables mitogenic stimulation while avoiding the risks associated with prolonged c-MYC overexpression.

In the current study, we noticed that beta cells transiently dedifferentiate during IVT mRNA-induced proliferation, as revealed by RT-qPCR and RNA-Seq analyses. These assays showed downregulation of key beta cell TFs, including Pdx1, Nkx6.1, and the maturation markers Ucn3 and Entpd3. These findings are consistent with previous studies, indicating that partial dedifferentiation from the mature phenotype facilitates replication of terminally differentiated cells such as beta cells and cardiomyocytes.^35,49–53^ In addition, the cell cycle progression assay showed that not all beta cells could complete the cell cycle. Potential explanations for cell cycle arrest include insufficient induction of regulators required for progression through the entire cell cycle, or individual cell-intrinsic limitations in cell cycle competency.^52,54–56^ Likewise, cellular senescence or checkpoint blockade due to insufficient or impaired DNA replication can also lead to cell cycle arrest.^12^

We acknowledge that our study has several limitations. First, we used dispersed islet cells rather than the intact pancreatic islets to assess the potential of IVT mRNA to stimulate beta-cell proliferation. IVT mRNA transfection is generally more efficient in monolayer cultures compared to compact cell clusters like pancreatic islets.^57^ Therefore, we anticipate that transfection efficiency and, consequently, beta-cell proliferation, would decrease if the IVT mRNAs were applied to intact islets. However, recent improvements in IVT mRNA transfection reagents and advances in delivery strategies could help overcome this issue.^57–60^ Second, we used human fetal pancreas-derived EndoC-BH5 cells to evaluate the potential of IVT mRNA to induce proliferation of human-derived beta cells. Although these cells are highly similar to native human beta cells in their functional and metabolic properties,^40^ EndoC-BH5 cells are unlikely to be as mature as adult pancreatic beta cells. As the maturity and differentiation state of beta cells significantly affect their replication potential,^3,49,61^ further research on mRNA-induced human beta-cell proliferation in the adult pancreatic islets is needed. Finally, although the stimulation of beta-cell proliferation is an attractive regenerative approach, unregulated beta-cell replication would have significant consequences. An extensive cell cycle activity occurred only during the first 3–4 days of mRNA-induced proliferation; thereafter, beta-cell proliferation attenuated to levels comparable to unstimulated control samples. This finding is consistent with IVT mRNA’s temporal character, which provides an important safety factor.

Nevertheless, further studies on the safety and applicability of IVT mRNA-induced proliferation are warranted to assess the relevance of our findings in the context of human adult beta-cell proliferation and its potential therapeutic application. If proven effective, IVT mRNA could serve as a tool for ex vivo expansion of donor islets before transplantation in diabetic patients. This strategy could improve the success of islet replacement therapies. In the future, targeted IVT mRNA delivery to residual beta cells in vivo, particularly in combination with immunomodulatory therapies, could support regeneration in autoimmune diabetes, paving the way for disease reversal.

In summary, our work demonstrates that IVT mRNA encoding cell cycle regulators can efficiently promote cell cycle activation and mitotic division in both rodent- and human-derived beta cells. These findings support the broader application of IVT mRNA technology in regenerative medicine. However, further validation in native human islets is needed before clinical application.

## MATERIALS AND METHODS

### Animals

We used 10–12-week-old Wistar male rats procured from (Charles River Laboratories, Germany). All animal experiments were approved by The Animal Care Committee of the Institute for Clinical and Experimental Medicine and Ministry of Health of Czech Republic (Permit Number: 30/2022; 53/2019). Animals were held according to the European Convention on Animal Protection and Guidelines on Research Animal Use in conventional breeding facility with 12/12 h light/dark cycle and free access to food and water.

### Islet isolation and culture

Rat islets were isolated from 4- to 6-month-old Wistar rats as previously described.^62^ Briefly, the pancreata of anesthetized rats were cannulated through the bile duct and filled with 15 mL of collagenase solution (Sigma-Aldrich, Germany, C9263). Collagenase-infused pancreata were then dissected and incubated at 37°C for 18 min with moderate shaking. Collagenase digestion was stopped using a cold Hank’s Balanced Salt Solution (HBSS) (Sigma-Aldrich, H8264) containing 1% heat-inactivated fetal bovine serum (FBS) (Sigma-Aldrich, F9665), before filtering the tissue suspension through a mesh. Pancreatic islets were then separated using a Ficoll (Sigma-Aldrich, F9878) discontinuous density gradient. After aspiration of the islet containing layer, the islet suspension was washed with HBSS, and the purified islets hand-picked under a stereomicroscope. Islets were then cultured in islet culture medium containing Connaught Medical Research Laboratories 1066 (CMRL-1066) medium (PanBiotech, Germany, P04-84600), 1mM GlutaMAX Supplement (Gibco, USA, 35050061), 10 mM HEPES Buffer Solution (Gibco, 15630049), and 10% FBS, at 37°C in a 5% CO2-buffered incubator.

### Islet cell culture

Isolated rat islets were collected and dispersed into a single-cell suspension by incubating for 20 min at room temperature (RT) in an Accutase solution (1 μL/islet; Sigma-Aldrich, A6964), followed by intense pipetting. Then, islet cells were washed once with HBSS, and seeded at 150,000 cells/cm^2^ density onto a decellularized matrix derived from the HTB-9 cell line (American Type Culture Collection, USA, 5637), in a 96-well plate (Greiner Bio-One, Germany, 655986). Islet cells were cultured in islet cell culture medium containing CMRL-1066 medium, 15% FBS, 1mM GlutaMAX Supplement, 1 mM Sodium Pyruvate (11360070), 1% Insulin-Transferrin-Selenium (41400045), and MEM Non-Essential Amino Acids Solution (11140035) (all from Gibco), at 37°C in a 5% CO2-buffered incubator until further use. For Click-IT EdU proliferation assays, an EdU analog (Invitrogen, A10044) was added to the culture medium after IVT mRNA transfection at a final concentration of 20 μM for the indicated time. The EdU containing islet cell culture medium was replaced every second day. For mitotic block induction, a CDK1 inhibitor RO-3306 (Sigma-Aldrich, SML0569) was added to the culture medium at a final concentration of 20 μM for the indicated time.

### EndoC-BH5 cell culture

EndoC-BH5 cells (Human Cell Design, France, BH5-WT-350) were cultured at 150,000 cells/cm^2^ on the decellularized matrix derived from the HTB-9 cell line in Ulti-B1 culture media (Human Cell Design, UB1-100-BSA), supplemented with 15% FBS, 1 mM GlutaMAX Supplement, 1 mM Sodium Pyruvate, 10 mM HEPES Buffer Solution, 1% Insulin-Transferrin-Selenium, MEM Non-Essential Amino Acids Solution, and 1% EmbryoMax Nucleosides (Sigma-Aldrich, ES-008) at 37°C in a 5% CO_2_-buffered incubator until further use. For Click-IT EdU proliferation assays, an EdU analog was added to the culture medium after IVT mRNA transfection for the indicated times, at a final concentration of 1 μM for flow cytometry analysis and 2 μM for immunofluorescence microscopy. The Ulti-B1 culture medium with all supplements was replaced every second day. For mitotic block induction, RO-3306 was added to the culture medium at a final concentration of 20 μM for the indicated time.

### Generation of IVT DNA templates

All sequences for IVT DNA templates are provided in Supplemental Information (Tables S1 to S12). Plasmids containing DNA template sequences were prepared by GeneArt gene synthesis service (Thermo Fisher Scientific, Germany). Plasmids were amplified in E. coli HST08 strain (Takara Bio, Japan, 636764) and purified using the CompactPrep Plasmid Midi Kit (Qiagen, Germany, 12843). Linearized plasmids were PCR amplified by the Q5 Hot Start High-Fidelity DNA polymerase (New England Biolabs, USA, M0493) and PCR products purified using the NucleoSpin Gel and PCR Clean-up Kit (Machery-Nagel, Germany, 740609.50). Sequence lengths were verified by 1.0% TBE-agarose gel electrophoresis and DNA template concentration were determined using the Qubit dsDNA BR Assay Kit (Invitrogen, USA, Q32850).

### *In vitro* mRNA transcription

In vitro transcription was performed with the MEGAscript T7 Transcription Kit (Invitrogen, AM1334) according to manufacturer’s instructions, using the PCR products of linearized plasmids encoding the DNA templates. 1-methylpseudouridine-5’-triphosphate (TriLink Biotechnologies, USA, N-1081) was used instead of uridine-5’-triphosphate to generate modified nucleoside-containing IVT mRNA and co-transcriptional capping of IVT mRNA was promoted by CleanCap Reagent AG (TriLink Biotechnologies, N-7113). After 3 h incubation at 37°C, the DNA template was digested with 1 μL Turbo DNase (Thermo Fisher Scientific, USA, AM2238) for 20 min at 37°C. Double-stranded RNA contaminants were removed using cellulose-based purification with cellulose fibers (Sigma-Aldrich, C6288) as previously described.^22^ To further reduce innate immune response, 5′-Triphosphate was removed by Antarctic Phosphatase (New England Biolabs, M0289S) according to manufacturer’s instructions. After final purification with MegaClear columns (Invitrogen, AM1908), the purified IVT mRNA was dissolved in RNAsecure Rnase Inactivation Reagent (Invitrogen, AM7005). RNA concentration was determined using a Qubit RNA BR Assay Kit (Invitrogen, Q10210), and purity and integrity were analyzed using Fragment Analyzer RNA Kit (Agilent, USA,DNF-471-0500) with a 5200 Fragment Analyzer System (Agilent). The prepared IVT mRNA was stored at −80°C until further use.

### IVT mRNA transfection

For transfection, the IVT mRNA was diluted in Opti-MEM (Invitrogen, 31985062) to a final concentration 40 ng/µL. Lipofectamine Messenger Max (Invitrogen, LMRNA001) was also diluted in Opti-MEM at a 1:24 ratio and incubated for 10 min at RT. To transfect cells in each well of a 96-well plate, 4 µL of diluted IVT mRNA (160 ng) was mixed with 10 µL diluted Lipofectamine Messenger Max and incubated for 8 min at RT. Then, the Lipofectamine/IVT mRNA complex was mixed with 40 µL of culture media and added to each well. When using combinations of various IVT mRNAs, equal amounts of each were used to a total 160 ng per well.

### Western blotting

Islet cells were lysed in RIPA Lysis Buffer (Sigma-Aldrich, R0278) with the complete Mini EDTA-free Protease Inhibitor Cocktail (Roche, Switzerland, 04693159001); protein concentration was measured with a BCA Protein Assay Kit (Thermo Fisher Scientific, 23227). A total of 20 µg of whole cell protein lysate diluted with 4X Laemmli loading buffer (250 mM Tris-HCl, pH 6.8; 8% SDS; 40% glycerol; 0.02% bromophenol blue; and 20% β-mercaptoethanol) was heated at 95°C for 3 min, loaded and separated on a 15% SDS-polyacrylamide gel, before transferring to a PVDF Western Blotting membrane (Roche, 3010040001). Then, membranes were blocked with 5% nonfat dried milk in TBS + 0.05% Tween, incubated with primary antibodies overnight at 4°C, washed and incubated with the corresponding horseradish peroxidase–conjugated secondary antibody for 1 h at RT. Finally, membranes were washed and imaged using a SuperSignal West Pico Plus Substrate (Thermo Fisher Scientific, 34577) and a G:BOX Chemi XR5 chemiluminescence detection system (Syngene, UK). All primary and secondary antibodies are provided in Table S13.

### Immunofluorescence and EdU staining

Cells grown on the decellularized matrix derived from the HTB-9 cell line were washed three-times with HBSS containing 1% FBS, and fixed with 4% paraformaldehyde (Polysciences, USA, 00380) in phosphate-buffered saline (PBS) for 20 min at RT. Cells were then permeabilized with 0.1% Triton X-100 in PBS for 20 min at RT, and blocked with blocking solution (5% donkey serum in PBS with 0.05% Triton X-100 and 0.75% Glycine) for 30 min at RT. For EdU staining, the Click-iT Plus EdU Cell Proliferation Kit (Invitrogen, C10640) was used before blocking and immunostaining according to manufacturer’s instructions. Then, cells were incubated with primary antibodies diluted in antibody diluent solution (0.2% cold fish skin gelatin, 0.01% sodium azide, 0.5% Triton X-100, 1% BSA in PBS) for 1 h at RT. Cells were washed three-times with PBS and incubated with the appropriate secondary antibodies diluted in blocking solution for 1 h at RT. Next, nuclei were stained with NucBlue Fixed Cell Stain ReadyProbes Reagent (Invitrogen, R37606) diluted in PBS for 5 min at RT and washed six-times with PBS before mounting with Dabco Mowiol solution (2.5% 1,4-diazabicyclo-[2,2,2]-octane, 4.8% Mowiol, 12% glycerol, and 0.2M Tris HCl). Immunofluorescent images were captured using an EVOS M7000 Imaging System (Invitrogen). All primary and secondary antibodies are provided in Table S13.

### Image analysis

For each sample, ≥ 30,000 cells were counted. Proliferation was calculated as the percentage of double-positive cells for EdU and each population marker with respect to the corresponding total cell population labeled with specific marker.

### Flow cytometry

Before flow cytometry, cells were detached from culture plates by incubation with pre-warmed TrypLE Select Enzyme (Gibco, A1217701) for 7 min at RT and further dissociated into a single-cell suspension using the Accutase solution for 20 min at RT. After intense pipetting and filtration through a 50 µm pore mesh, cells were washed twice with ice-cold wash buffer (1% BSA in PBS) and incubated with Fixable Viability Dye eFluor 450 (Invitrogen, 65-0863-14) staining solution (0.1% in PBS) for 30 min at 4°C. Then, cells were washed twice with wash buffer, fixed with 4% formaldehyde in PBS for 20 min at 4°C, and permeabilized with Permeabilization Buffer (Invitrogen, 00-8333-56) for 15 min at RT. For EdU analysis, the Click-iT Plus EdU Cell Proliferation Kit (Invitrogen, C10640) was used according to manufacturer’s instructions. Briefly, fixed and permeabilized cells were incubated in the Click-iT reaction cocktail for 30 min at RT in the dark before washing three-times with Permeabilization Buffer.

For subsequent insulin staining, cells were resuspended and incubated in blocking buffer (2% rat and 2% donkey serum in Permeabilization Buffer) for 15 min at RT. After blocking, cells were incubated with the primary rabbit anti-insulin antibody (Abcam, UK, ab181547) in a 1:400 dilution for 60 min at RT, washed three-times with Permeabilization Buffer, and further incubated with the secondary donkey anti-Rabbit IgG Alexa Fluor 555 Secondary Antibody (Invitrogen, A-31572) at 1:100 for 30 min at RT.

Finally, cells were washed three-times with Permeabilization Buffer and analyzed on a BD LSR II flow cytometer (BD Biosciences, USA). In addition to Forward Scatter (FSC) and Side Scatter (SSC)parameters, signals for AlexaFluor 555 (561 nm laser: 586/15), AlexaFluor 647 (637 nm laser: 660/20), GFP (488 nm laser: 525/50), and FVD eFluor 450 (405 nm laser: 450/50) were acquired. Data were analyzed using FlowJo v10.8.1 software (BD Biosciences). EdU-positive beta-cells were quantified as the percentage of the total number of analyzed beta-cells. At least 20,000 events were acquired per sample.

### Flow cytometry-based cell cycle progression assay

For this assay, cells were harvested and processed as for standard flow cytometry until fixation. In addition to standard fixation with 4% formaldehyde in PBS, an additional fixation step with ice-cold 70% ethanol for 20 min at −20°C was performed. After fixation, cells were washed three-times with ice-cold wash buffer (1% BSA in PBS) and permeabilized with Permeabilization Buffer (Invitrogen, 00-8333-56) for 15 min at RT. For EdU analysis, the Click-iT Plus EdU Cell Proliferation Kit (Invitrogen, C10637) was used according to manufacturer’s instructions. Briefly, fixed and permeabilized cells were incubated in the Click-iT reaction cocktail for 30 min at RT in the dark, before washing three-times with Permeabilization Buffer. For subsequent insulin staining, cells were resuspended and incubated in blocking buffer (2% rat and 2% donkey serum in Permeabilization Buffer) for 15 min at RT. After blocking, EndoC-BH5 cells were incubated with the primary rabbit anti-insulin antibody (Abcam, ab181547) at a 1:400 dilution for 60 min at RT, washed three-times with Permeabilization Buffer, and further incubated with a secondary donkey anti-Rabbit IgG Alexa Fluor 555 Secondary Antibody (Invitrogen, A-31572) at a 1:100 dilution, for 30 min at RT. Then, rat islet cells were incubated with the primary mouse anti-insulin antibody (Sigma-Aldrich, I2018) at a 1:200 dilution for 60 min at RT, washed three-times with Permeabilization Buffer, and further incubated with a secondary rat anti-Mouse IgG BV711 Secondary Antibody (BD Biosciences, 565786) at a 1:100 dilution for 30 min at RT.

Finally, cells were washed three-times with Permeabilization Buffer, DNA was stained with Hoechst33342 at a 1:4000 dilution (Invitrogen, C10637G) and analyzed with a FACSymphony (BD Biosciences) flow cytometer. In addition to FSC and SSC parameters, signals for Hoechst33342 (355nm, laser: 450/50), AlexaFluor488 (488nm laser: 530/30), AlexaFluor 555 (561nm laser: 586/15), AlexaFluor 647 (637nm laser: 670/30), and BV711 (405nm laser: 710/50) were acquired. Data were analyzed using FlowJo v10.8.1 software (BD Biosciences).

### Beta-cell sorting and quantification

Before beta-cell sorting, isolated rat islets were collected and dispersed into a single-cell suspension by incubation in the Accutase solution (1 μL/islet; Sigma-Aldrich, A6964) for 20 min at RT, followed by intense, continuous pipetting. For beta-cell staining, dispersed islet cells were centrifuged at 200 g for 3 min, resuspended in islet culture medium (10% FBS and 1 mM GlutaMAX supplement in CMRL-1066 medium), and incubated with 1 μM FluoZin-3, AM (Invitrogen, F24195), previously diluted with Pluronic F-127 (Invitrogen, P3000MP), at 37°C for 30 min in the dark. Then, cells were washed twice with islet culture medium and incubated in the same medium for 30 min at 37°C in the dark. Finally, cells were centrifuged at 200 g for 3 min, resuspended in islet culture medium containing 2% FBS, filtered through a 50 µm pore mesh, and sorted using a BD FACSMelody™ Cell Sorter (BD Biosciences) with a 488 nm excitation laser coupled with a 527/32 nm filter. A total of 30,000 FluoZin-3-positive cells/well were sorted and seeded directly into 96-well plates, precoated with the decellularized matrix derived from HTB-9 cell line and prefilled with 100 μL FBS. At 8 h after cell sorting and attachment to the culture surface, the sorting solution was replaced by islet cell culture medium. Two days after sorting, transfection with the stimulatory IVT mRNA (CCND1 T283A + CDK4) was performed, with addition of 20 µM EdU analog. The islet cell culture medium containing EdU was replaced every second day. The EdU proliferation assay and immunostaining for insulin and DAPI was performed 4 days after IVT mRNA transfection, as described in the Immunofluorescence and EdU staining section. Stitched images covering the entire surface of each well were captured using the EVOS M7000 Imaging System (Invitrogen). To quantify the purity and total number of beta cells in each well, insulin- and DAPI-positive cells were counted in stitched images using Celleste Image Analysis Software (Invitrogen). In addition, the proportion of EdU+ beta cells was determined in IVT mRNA-treated and untreated control samples. The purity of the sorted beta cells was > 99.0%, as confirmed by insulin immunostaining of the sorted cells.

### RNA isolation and RT-qPCR

RNA was isolated from cultured islet cells using an RNeasy Mini Plus kit (Qiagen, 74134) according to manufacturer’s instructions. cDNA was prepared using the Transcriptor First Strand cDNA Synthesis Kit (Roche, 11483188001). qPCRs were performed using the FastStart Universal SYBR Green Master Rox (Roche, 4913914001) with gene-specific primers non-amplifying IVT mRNA constructs (Integrated DNA Technologies, Netherlands). Primer sequences are provided in table S14. Fold changes in gene expression were determined using the ΔΔCT method, with normalization according to *Hprt* and *Actb* expression, as previously reported.^62^

### Bulk-cell RNA sequencing and analyses

For the bulk-cell RNA sequencing protocol, total RNA was used as input to prepare sequencing libraries using the QuantSeq 3′ mRNA-Seq Library Prep Kit FWD for Illumina (Lexogen), with the UMI Second Stranded Synthesis Module for QuantSeq FWD following the manufacturer’s directions. RNA-Seq libraries were prepared from FACS purified beta-cell samples treated with CCND1 T286A and CDK4; GFP IVT mRNA; and untreated controls, each comprising five biological replicates per condition. Quality control included quantification using the Qubit dsDNA assay kit (Invitrogen, Q32851) and Qubit 4 fluorometer, and fragment assessment using a Fragment Analyzer System with the NGS Fragment Kit (Agilent). Libraries were sequenced on a NextSeq 500 instrument using the NextSeq 500/550 High Output Kit v2.5 75 Cycles (Illumina, USA, 200024906) in 1 × 76 bp mode, generating an average of 23,5 million reads per sample. Raw read quality was checked with FastQC v0.11.9 and contamination with FastQ_Screen v0.11.1. Then, the 6 bp-long UMIs followed by “TATA” spacer were added to reads name using umi_tools extract function (https://genome.cshlp.org/content/early/2017/01/18/gr.209601.116.abstract). Subsequently, adaptor and low-quality read trimming were performed by TrimmomaticSE v0.36 using specified parameters, and ribosomal/mitochondrial reads were filtered with SortMeRNA v2.1b. Reads were aligned to the Rattus norvegicus genome (Rnor 6.0) using STAR v2.5.2b, converted to BAM files. The final gene count tables were generated using htseq-count v0.11.3 against Rnor.6.0.93.gtf. Reads for *Ccnd1, Ccnd2, Ccnd3, Cdk4, Cdk6* and *Ins2* were excluded due to a significant homology with the IVT mRNA sequences used for transfections. All bioinformatic analyses were conducted in R (v4.2.2), converting ENSEMBL IDs to gene symbols using the org.Rn.eg.db database. Differential gene expression (DGE) analysis was executed with the DESeq2 package (v1.36). Differentially expressed genes (DEGs) were stringently identified by the criteria of an adjusted P-value (Padj < 0.05) and an absolute log2fold change |log2FC| > 1. Finally, PCA and heatmaps were generated from rlog-transformed data, and Gene Ontology (GO) analysis was performed using the clusterProfiler package (v4.4.4).

### Statistical analysis

All studies in rodent and human cells were performed in ≥ 3 independent sets of experiments. The unpaired two-sided students’ t-test was used to analyze two groups’ samples. An analysis of variance (ANOVA) with appropriate corrections for post hoc analysis (Bonferroni or Tukey) was used for multiple group comparisons. Unless otherwise stated, data are expressed as mean ± standard deviation (SD). Results were considered significant for a P-value ≤ 0.05. Exact p-values are labeled in each figure. GraphPad Prism V10 (GraphPad Software Inc.) was used for statistical analysis. All graphs were generated using RStudio (version 2026.07.1 Pacific Dogwood) running R (version 4.6.1 Happy Hop). Specific plots were constructed using the packages ggplot2 (v4.0.3) and ggpubr (v1.0.0). For RNA sequencing analysis, a Benjamini-Hochberg correction was used to adjust for multiple testing, with an adjusted P-value < 0.05 (i.e., false discovery rate < 10%). For GO analysis, only pathways with a P-value < 0.05 were reported.

## DATA AVAILABILITY

All data needed to evaluate the conclusions in the paper are present in the paper or the supplementary materials. Additional data related to this paper may be requested from the authors. RNA-seq datasets have been deposited in the Gene Expression Omnibus under accession number GSE312170.

### Lead contact

- Requests for further information and resources should be directed to and will be fulfilled by the lead contact, Tomas Koblas.

## ACKNOWLEDGMENTS

This research was supported by the project National Institute for Research of Metabolic and Cardiovascular Diseases (Programme EXCELES, ID Project No. LX22NPO5104) - Funded by the European Union–Next Generation EU (J.K. and FS). Additional support was provided by the: Czech Science Foundation, project number 24-11364S/26-11364S (T.K. and L.V.); Ministry of Health, Czech Republic - conceptual development of research organization („Institute for Clinical and Experimental Medicine–IKEM, IN 00023001“) (F.S.); Institutional support RVO 86652036 (L.V.); Charles University, project GA UK No. 299122 (K.B.); and NEURON Foundation, project 20/2017 (T.K.). We thank Mikhail Okun from Thermo Fisher Scientific for developing the Celleste software script used for High-content image analysis, Michal Koblas for creating R scripts for graphs generation, and Ivan Leontovyc and Magdalena Spitalnikova Vertatova for assistance with Western blotting and beta-cell FACS sorting. We also thank the Šuran family for providing additional financial support.

## AUTHOR CONTRIBUTIONS

Conceptualization of the project, T.K.; methodology, T.K., K.B., K.Z., P.A., and P.G.; investigation, T.K., K.B., K.Z., and P.A.; visualization, T.K., K.B., and P.A.; writing – original draft, T.K.; writing – review & editing, T.K., K.B., L.V., J.K., and F.S.; funding acquisition, T.K., K.B., L.V., J.K., and F.S.; supervision, T.K., L.V., P.G, and F.S. All authors reviewed and approved the final version of the manuscript.

## DECLARATION OF INTERESTS

All other authors declare they have no competing interests.

## DECLARATION OF GENERATIVE AI AND AI-ASSISTED TECHNOLOGIES IN THE WRITING PROCESS

The AI tools were used only to improve the grammar of the text. Generative AI was used to prepare the beta cell icons of the graphical abstract in BioRender.

## SUPPLEMENTAL INFORMATION

Document S1. Supplemental Figures S1–S7 and Tables S1–S14.

## Notes

### Competing Interest Statement

The authors have declared no competing interest.

