## SUPPLEMENTAL INFORMATION for "Stimulation of rodent and human beta-cell proliferation using synthetic modified mRNAs encoding cell cycle regulators"

This PDF file includes:

Figures S1 to S7 and Tables S1 to S14

**Supplemental Figures**

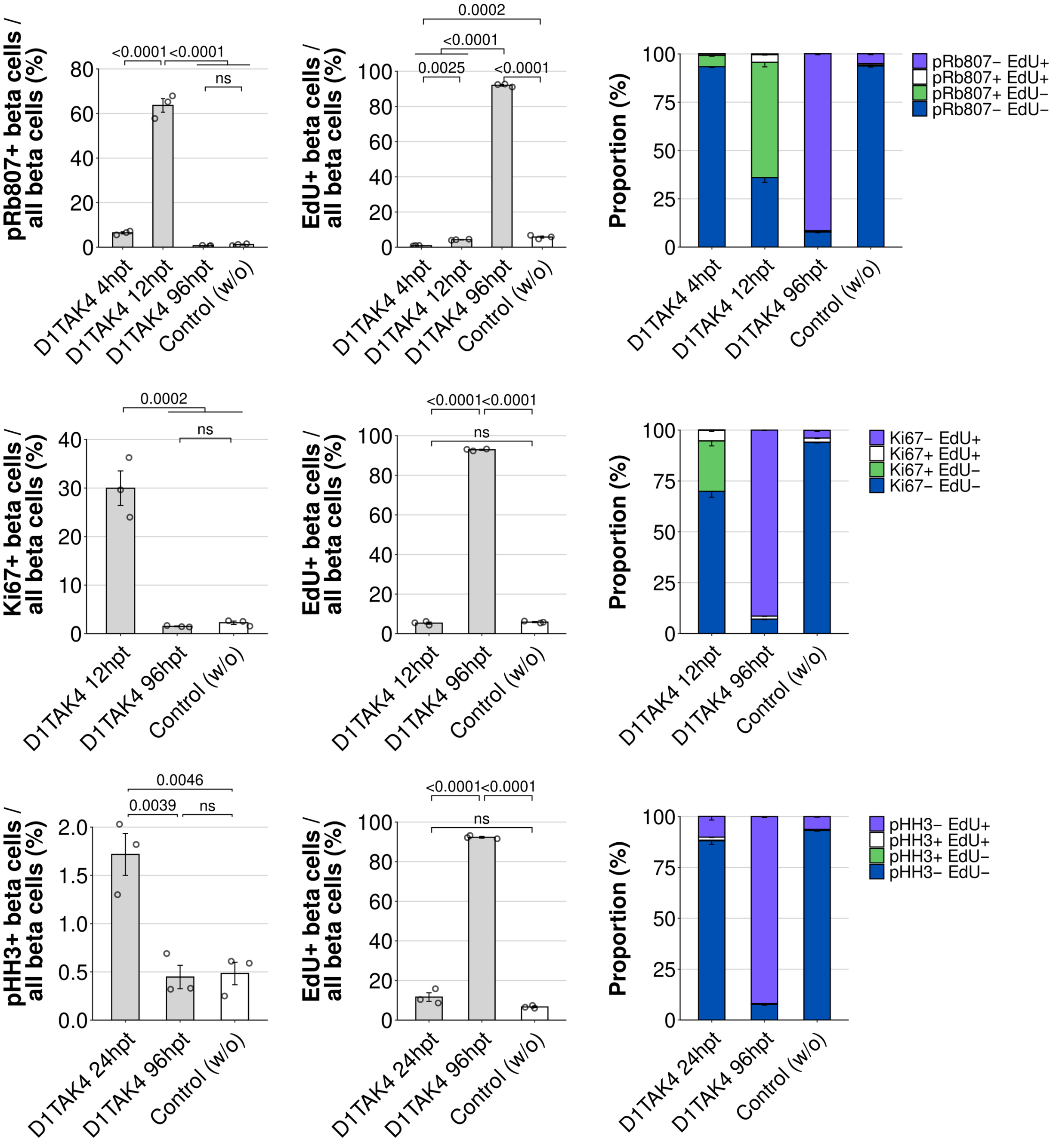


Figure S1. Analysis of cell cycle kinetics during the IVT mRNA induced beta-cell proliferation. High-content imaging-based quantification of pRb807+ (A); Ki67+ (B); pHH3+ (C); and EdU+ (A, B, C) beta cells (left and middle), and the corresponding proportions of beta-cell subpopulations based on specific cell cycle marker/EdU positivity (right), in CCND1T286A/CDK4 IVT mRNA-treated (abbreviated as D1TAK4) islet cell samples, and control untreated samples, at specified time points post transfection. Data represent mean ± SD (n = 3 biologically independent samples per group). Statistical significance was determined using one-way ANOVA with Bonferroni's post-test analysis. P-values are shown above the graphs.

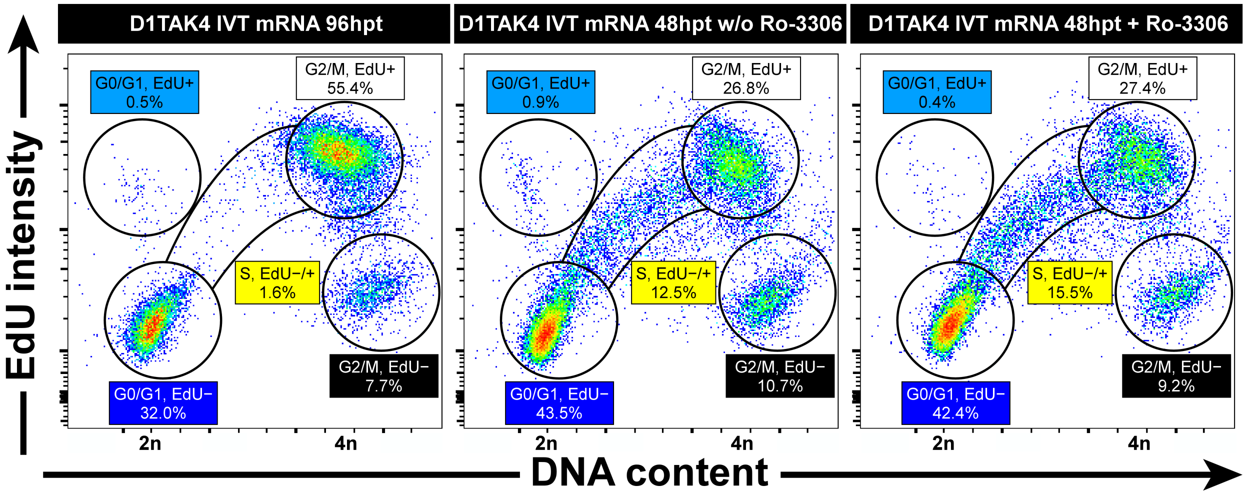


Figure S2. IVT mRNAs encoding cyclin D1 and CDK4 only activates cell cycle entry and progression, not mitosis, in human fetal beta-cell-derived EndoC-BH5 cells.

Representative flow cytometry plots of cell cycle subpopulations in CCND1T286A and CDK4 IVT mRNA-treated EndoC-BH5 cells (96 hpt and 48hpt); and verification of the cell cycle progression assay using CDK1 inhibitor Ro-3306 in IVT mRNA-treated (48 hpt) EndoC-BH5 cells. Circles and S-phase corridor indicate the gated populations.

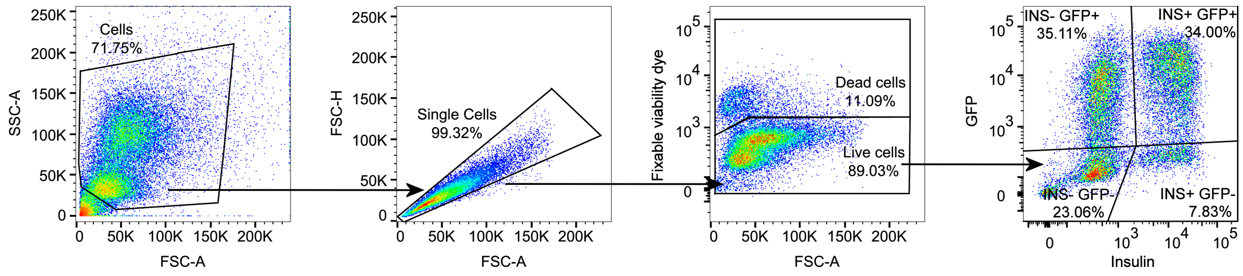

Figure S3. Gating strategy for flow cytometry analysis of GFP IVT mRNA transfection efficiency. The gating strategy corresponds to the flow cytometry plots presented in Figure 1E. Cells were gated to identify singlets (SSC-A vs. FSC-A; FSC-H vs. FSC-A), live cells (Fixable viability dye vs. FSC-A), and GFP-positive beta-cells and non-beta cells (GFP vs. insulin).

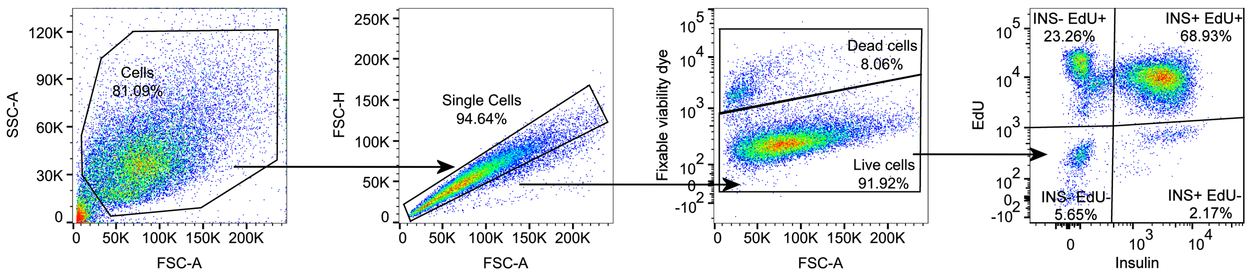


Figure S4. Gating strategy for flow cytometry analysis of EdU incorporation in cyclins D/CDKs IVT mRNA-treated islet cell samples. The gating strategy corresponds to the flow cytometry plots presented in Figure 2C. Cells were gated to identify singlets (SSC-A vs. FSC-A; FSC-H vs. FSC-A), live cells (Fixable viability dye vs. FSC-A), and EdU-positive beta-cells and non-beta cells (EdU vs. insulin).

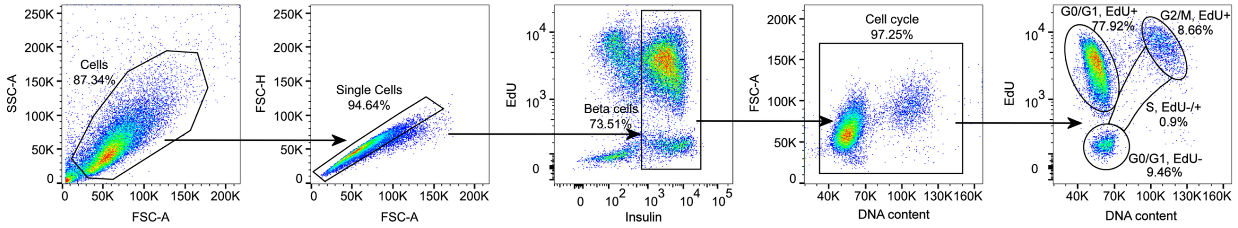


Figure S5. Gating strategy for flow cytometry analysis of cell cycle progression in rat islet cells. The gating strategy corresponds to the flow cytometry plots presented in Figures 4D and 4E. Cells were gated to identify singlets (SSC-A vs. FSC-A; FSC-H vs. FSC-A), beta cells (EdU vs. insulin), proliferating and quiescent beta-cells (FSC-A vs. DNA amount), and beta-cells in specific cell-cycle phases (EdU vs. DNA amount).

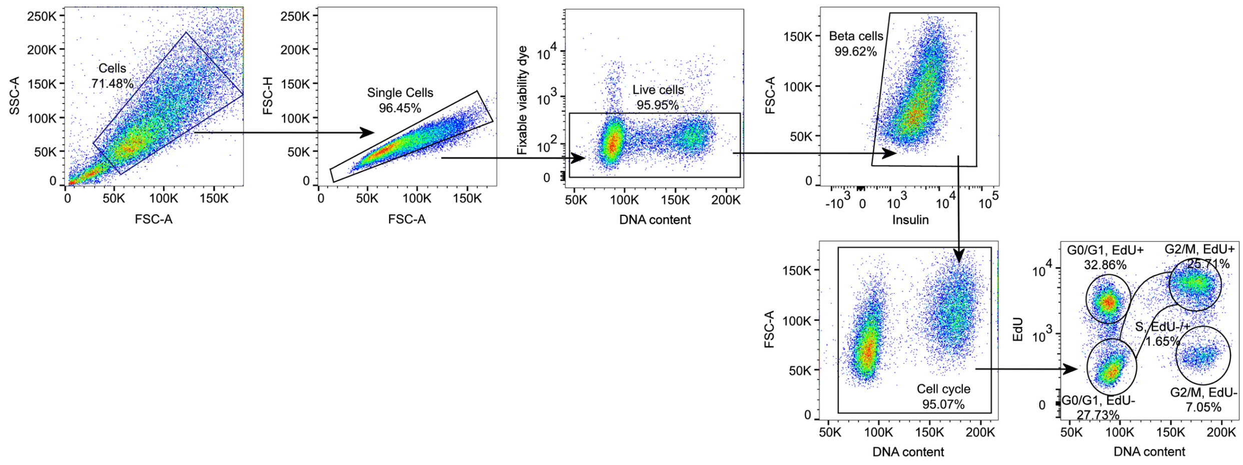


Figure S6. Gating strategy for flow cytometry analysis of cell cycle progression in human fetal beta-cell derived EndoC-BH5 cells.

The gating strategy corresponds to the flow cytometry plots presented in Figures 7C and 7D. Cells were gated to identify singlets (SSC-A vs. FSC-A; FSC-H vs. FSC-A), live cells (Fixable viability dye vs. FSC-A), beta cells (FSC-A vs. insulin), proliferating and quiescent beta-cells (FSC-A vs. DNA amount), and beta-cells in specific cell-cycle phases (EdU vs. DNA amount).

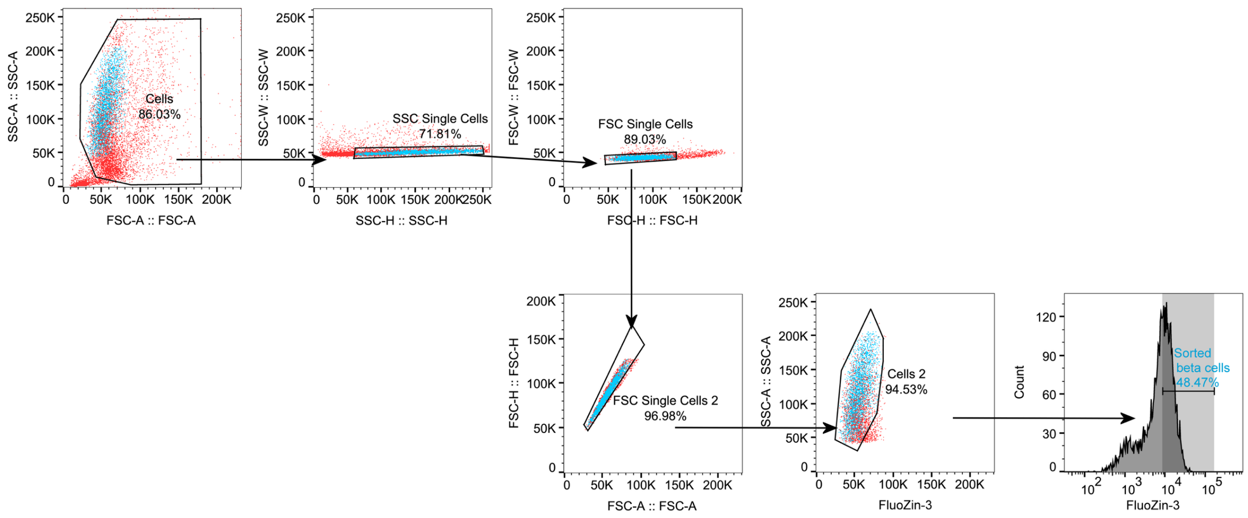


Figure S7. Gating strategy for FACS sorting rat beta-cells.

The gating strategy corresponds to the High-content imaging-based analysis of FACS-sorted beta-cells in Figures 4A and 4B; and RNA-seq analysis of FACS-sorted beta-cells in Figure 6. Cells were gated to identify singlets (SSC-A vs. FSC-A; SSC-W vs. SSC-H; FSC-W vs. FSC-H; FSC-H vs. FSC-A), and purified beta-cells (SSC-A vs. Fluozin-3, and Fluozin-3 intensity). Beta cells are shown in blue in backgating.

### Supplemental Tables

Table S1. DNA template sequence encoding GFP IVT mRNA.

| Element | Description | Position |
|---|---|---|
| T7 promoter | T7 RNA polymerase promoter sequence | 1-17 |
| Cap | A modified 5'-cap1 structure ($m^7G$-5'-ppp-5'-$Am^{2'}$) | 18-19 |
| 5'-UTR | 5´-untranslated region derived from rat insulin2 mRNA (NM_019130.2) with an optimized Kozak sequence | 20-77 |
| AcGFP1 | Sequence encoding AcGFP1 (in bold), constitutively fluorescent green fluorescent protein derived from Aequorea coerulescens; stop codons: 795-800 (underlined) | 78-800 |
| 3'-UTR | The 3´-untranslated region derived from rat insulin2 mRNA (NM_019130.2) | 801-858 |
| poly(A) | A 110-nucleotide poly(A)-tail consisting of a stretch of 30 adenosine residues, followed by a 10-nucleotide linker sequence and another 70 adenosine residues. | 859-968 |

Sequence:

TAATACGACTCACTATAAGGAGCCCTAAGTGACCAGCTACAGTCGGAAAC **50**
CATCAGCAAGCAGGTCATTGTGCCAAC**ATGGTGAGCAAGGGCGCCGAGCT** **100**
**GTTCACCGGCATCGTGCCCATCCTGATCGAGCTGAATGGCGATGTGAATG** **150**
**GCCACAAGTTCAGCGTGAGCGGCGAGGGCGAGGGCGATGCCACCTACGGC** **200**
**AAGCTGACCCTGAAGTTCATCTGCACCACCGGCAAGCTGCCTGTGCCCTG** **250**
**GCCCACCCTGGTGACCACCCTGAGCTACGGCGTGCAGTGCTTCTCACGCT** **300**
**ACCCCGATCACATGAAGCAGCACGACTTCTTCAAGAGCGCCATGCCTGAG** **350**
**GGCTACATCCAGGAGCGCACCATCTTCTTCGAGGATGACGGCAACTACAA** **400**
**GTCGCGCGCCGAGGTGAAGTTCGAGGGCGATACCCTGGTGAATCGCATCG** **450**
**AGCTGACCGGCACCGATTTCAAGGAGGATGGCAACATCCTGGGCAATAAG** **500**
**ATGGAGTACAACTACAACGCCCACAATGTGTACATCATGACCGACAAGGC** **550**
**CAAGAATGGCATCAAGGTGAACTTCAAGATCCGCCACAACATCGAGGATG** **600**
**GCAGCGTGCAGCTGGCCGACCACTACCAGCAGAATACCCCCATCGGCGAT** **650**
**GGCCCTGTGCTGCTGCCCGATAACCACTACCTGTCCACCCAGAGCGCCCT** **700**
**GTCCAAGGACCCCAACGAGAAGCGCGATCACATGATCTACTTCGGCTTCG** **750**
**TGACCGCCGCCGCCATCACCCACGGCATGGATGAGCTGTACAAG**TGATAA **800**
GCCCACCACTACCCTGTCCACCCCTCTGCAATGAATAAAACCTTTGAAAG **850**
AGCACTACAAAAAAAAAAAAAAAAAAAAAAAAAAAAAAGCATATGACTAA **900**
AAAAAAAAAAAAAAAAAAAAAAAAAAAAAAAAAAAAAAAAAAAAAAAAAA **950**
AAAAAAAAAAAAAAAAAA **968**

Table S2. DNA template sequence encoding rat Ccnd1 IVT mRNA.

| Element | Description | Position |
| --- | --- | --- |
| T7 promoter | T7 RNA polymerase promoter sequence | 1-17 |
| Cap | A modified 5’-cap1 structure ($m^7G$-5'-ppp-5'-$Am^{2'}$) | 18-19 |
| 5’-UTR | 5´-untranslated region derived from rat insulin2 mRNA (NM_019130.2) with an optimized Kozak sequence | 20-77 |
| Ccnd1 | Sequence encoding rat Ccnd1 (NM_171992.6; in bold); stop codons: 963-968 (underlined) | 78-968 |
| 3’-UTR | The 3´-untranslated region derived from rat insulin2 mRNA (NM_019130.2) | 969-1026 |
| poly(A) | A 110-nucleotide poly(A)-tail consisting of a stretch of 30 adenosine residues, followed by a 10-nucleotide linker sequence and another 70 adenosine residues. | 1027-1136 |

Sequence:

TAATACGACTCACTATAAGGAGCCCTAAGTGACCAGCTACAGTCGGAAAC **50**
CATCAGCAAGCAGGTCATTGTGCCAAC**ATGGAACACCAGCTCCTGTGCTG** **100**
**CGAAGTGGAGACCATCCGCCGCGTACCCTGACACCAATCTCCTCAACG** **150**
**ACCGGGTGCTGCGAGCCATGCTTAAGACTGAGGAGACCTGCGCGCCCTCC** **200**
**GTTTCTTACTTCAAGTGCGTGCAGAGGGAGATTGTGCCATCCATGCGGAA** **250**
**AATCGTGGCCACCTGGATGCTAGAGGTCTGCGAGGAGCAGAAGTGCGAAG** **300**
**AGGAGGTCTTCCCGCTGGCCATGAACTACCTGGACCGTTTCTTGTCTCTG** **350**
**GAGCCCCTGAAGAAGAGCCGCCTGCAGCTTCTGGGGGCCACCTGCATGTT** **400**
**CGTGGCCTCTAAGATGAAGGAGACCATTCCCCTGACTGCCGAGAAGTTGT** **450**
**GCATCTACACTGACAACTCTATCCGCCCCGAGGAGTTGCTGCAAATGGAA** **500**
**CTGCTTCTGGTGAACAAGCTCAAATGGAACCTGGCCGCCATGACTCCCCA** **550**
**CGATTTCATCGAACACTTCCTCTCCAAAATGCCAGAGGCGGATGAGAACA** **600**
**AGCAGATCATCCGCAAACATGCACAGACCTTTGTGGCCCTCTGTGCCACA** **650**
**GATGTGAAGTTCATTTCCAACCCACCCTCCATGGTGGCTGCTGGGAGTGT** **700**
**GGTGGCCGCGATGCAGGGCCTGAACCTGGGCAGCCCCAACAACTTCCTCT** **750**
**CCTGCTACCGCACAACGCACTTTCTTTCCAGAGTCATCAAGTGTGACCCG** **800**
**GACTGCCTCCGTGCCTGCCAGGAACAGATTGAAGCCCTTCTGGAGTCAAG** **850**
**CCTGCGCCAGGCCCAGCAGAACATCGATCCCAAGGCCACCGAGGAGGAAG** **900**
**GGGAAGTGGAGGAAGAAGCTGGTCTGGCTTGCACACCCACCGACGTCCGA** **950**
**GATGTGGACATC**TGATAAGCCCACCACTACCCTGTCCACCCCTCTGCAAT **1000**
GAATAAAACCTTTGAAAGAGCACTACAAAAAAAAAAAAAAAAAAAAAAAA **1050**
AAAAAAGCATATGACTAAAAAAAAAAAAAAAAAAAAAAAAAAAAAAAAAA **1100**
AAAAAAAAAAAAAAAAAAAAAAAAAAAAAAAAAAAAA **1137**

Table S3. DNA template sequence encoding rat Ccnd1 T286A IVT mRNA.

| **Element** | **Description** | **Position** |
|---|---|---|
| T7 promoter | T7 RNA polymerase promoter sequence | 1-17 |
| Cap | A modified 5'-cap1 structure ($m^7G$-5'-ppp-5'-$Am^{2'}$) | 18-19 |
| 5'-UTR | 5´-untranslated region derived from rat insulin2 mRNA (NM_019130.2) with an optimized Kozak sequence | 20-77 |
| Ccnd1 T286A | Sequence encoding rat Ccnd1 (NM_171992.6; in bold), with introduced mutation T286A (threonine to alanine at position 286); stop codons: 963-968 (underlined) | 78-968 |
| 3'-UTR | The 3´-untranslated region derived from rat insulin2 mRNA (NM_019130.2) | 969-1026 |
| poly(A) | A 110-nucleotide poly(A)-tail consisting of a stretch of 30 adenosine residues, followed by a 10-nucleotide linker sequence and another 70 adenosine residues. | 1027-1136 |

Sequence:

| | |
|---|---|
| TAATACGACTCACTATAAGGAGCCCTAAGTGACCAGCTACAGTCGGAAAC | **50** |
| CATCAGCAAGCAGGTCATTGTGCCAAC**ATGGAACACCAGCTCCTGTGCTG** | **100** |
| **CGAAGTGGAGACCATCCGCCGCGTACCCTGACACCAATCTCCTCAACG** | **150** |
| **ACCGGGTGCTGCGAGCCATGCTTAAGACTGAGGAGACCTGCGCGCCCTCC** | **200** |
| **GTTTCTTACTTCAAGTGCGTGCAGAGGGAGATTGTGCCATCCATGCGGAA** | **250** |
| **AATCGTGGCCACCTGGATGCTAGAGGTCTGCGAGGAGCAGAAGTGCGAAG** | **300** |
| **AGGAGGTCTTCCCGCTGGCCATGAACTACCTGGACCGTTTCTTGTCTCTG** | **350** |
| **GAGCCCCTGAAGAAGAGCCGCCTGCAGCTTCTGGGGGCCACCTGCATGTT** | **400** |
| **CGTGGCCTCTAAGATGAAGGAGACCATTCCCCTGACTGCCGAGAAGTTGT** | **450** |
| **GCATCTACACTGACAACTCTATCCGCCCCGAGGAGTTGCTGCAAATGGAA** | **500** |
| **CTGCTTCTGGTGAACAAGCTCAAATGGAACCTGGCCGCCATGACTCCCCA** | **550** |
| **CGATTTCATCGAACACTTCCTCTCCAAAATGCCAGAGGCGGATGAGAACA** | **600** |
| **AGCAGATCATCCGCAAACATGCACAGACCTTTGTGGCCCTCTGTGCCACA** | **650** |
| **GATGTGAAGTTCATTTCCAACCCACCCTCCATGGTGGCTGCTGGGAGTGT** | **700** |
| **GGTGGCCGCGATGCAGGGCCTGAACCTGGGCAGCCCCAACAACTTCCTCT** | **750** |
| **CCTGCTACCGCACAACGCACTTTCTTTCCAGAGTCATCAAGTGTGACCCG** | **800** |
| **GACTGCCTCCGTGCCTGCCAGGAACAGATTGAAGCCCTTCTGGAGTCAAG** | **850** |
| **CCTGCGCCAGGCCCAGCAGAACATCGATCCCAAGGCCACCGAGGAGGAAG** | **900** |
| **GGGAAGTGGAGGAAGAAGCTGGTCTGGCTTGCGCACCCACCGACGTCCGA** | **950** |
| **GATGTGGACATC**TGATAAGCCCACCACTACCCTGTCCACCCCTCTGCAAT | **1000** |
| GAATAAAACCTTTGAAAGAGCACTACAAAAAAAAAAAAAAAAAAAAAAAA | **1050** |
| AAAAAAGCATATGACTAAAAAAAAAAAAAAAAAAAAAAAAAAAAAAAAAA | **1100** |
| AAAAAAAAAAAAAAAAAAAAAAAAAAAAAAAAAAAA | **1137** |

Table S4. DNA template sequence encoding rat Ccnd2 IVT mRNA.

| Element | Description | Position |
|---|---|---|
| T7 promoter | T7 RNA polymerase promoter sequence | 1-17 |
| Cap | A modified 5’-cap1 structure ($m^7G$-5'-ppp-5'-$Am^{2'}$) | 18-19 |
| 5’-UTR | 5´-untranslated region derived from rat insulin2 mRNA (NM_019130.2) with an optimized Kozak sequence | 20-77 |
| Ccnd2 | Sequence encoding rat Ccnd2 (NM_022267.2; in bold); stop codons: 942-947 (underlined) | 78-947 |
| 3’-UTR | The 3´-untranslated region derived from rat insulin2 mRNA (NM_019130.2) | 948-1005 |
| poly(A) | A 110-nucleotide poly(A)-tail consisting of a stretch of 30 adenosine residues, followed by a 10-nucleotide linker sequence and another 70 adenosine residues. | 1006-1115 |

Sequence:

| | |
|---|---|
| TAATACGACTCACTATAAGGAGCCCTAAGTGACCAGCTACAGTCGGAAAC | **50** |
| CATCAGCAAGCAGGTCATTGTGCCAAC**ATGGAGCTGCTGTGCTGTGAGGT** | **100** |
| **GGACCCGGTCCGCAGGGCCGTGCCGGACCGCAACCTGCTGGAAGACCGCG** | **150** |
| **TCCTGCAGAACCTGTTGACTATCGAGGAGCGCTACCTCCCGCAGTGTTCC** | **200** |
| **TATTTCAAGTGCGTGCAGAAGGACATCCAGCCGTACATGCGCAGGATGGT** | **250** |
| **GGCTACCTGGATGCTAGAGGTCTGTGAGGAACAGAAGTGTGAAGAAGAGG** | **300** |
| **TCTTTCCTCTGGCCATGAATTACCTGGACCGTTTCTTGGCTGGAGTCCCG** | **350** |
| **ACTCCTAAGACCCATCTCCAGCTCCTGGGCGCTGTGTGCATGTTCCTAGC** | **400** |
| **TTCCAAGCTGAAAGAGACCATCCCGCTGACTGCCGAAAAGCTGTGTATTT** | **450** |
| **ACACCGACAATTCTGTGAAACCCCAGGAGCTGCTGGAGTGGGAACTGGTG** | **500** |
| **GTGCTGGGTAAGCTGAAGTGGAACCTGGCTGCAGTAACCCCTCACGACTT** | **550** |
| **CATTGAGCACATCCTACGCAAGCTGCCCCAGCAGAAGGAGAAGCTGTCCC** | **600** |
| **TGATCCGCAAGCATGCGCAGACCTTCATCGCTCTGTGTGCTACCGACTTC** | **650** |
| **AAGTTTGCCATGTACCCGCCATCGATGATCGCAACTGGAAGCGTGGGAGC** | **700** |
| **AGCCATCTGCGGGCTTCAGCAGGACGAGGAAGTGAATGCACTCACGTGCG** | **750** |
| **ATGCCCTGACGGAGCTGCTGGCCAAGATCACCCACACCGATGTGGATTGT** | **800** |
| **CTCAAAGCCTGCCAGGAGCAAATCGAGGCTGTGCTGCTTAACAGCCTTCA** | **850** |
| **GCAGTTCCGTCAAGAGCAGCACAACGGCTCCAAGTCTGTGGAAGATCCGG** | **900** |
| **ACCAAGCCACCACCCCTACAGACGTGCGGGATGTTGACCTG**TGATAAGCC | **950** |
| CACCACTACCCTGTCCACCCCTCTGCAATGAATAAAACCTTTGAAAGAGC | **1000** |
| ACTACAAAAAAAAAAAAAAAAAAAAAAAAAAAAAAGCATATGACTAAAAA | **1050** |
| AAAAAAAAAAAAAAAAAAAAAAAAAAAAAAAAAAAAAAAAAAAAAAAAAA | **1100** |
| AAAAAAAAAAAAAAA | **1115** |

Table S5. DNA template sequence encoding rat Ccnd2 T280A IVT mRNA.

| Element | Description | Position |
|---|---|---|
| T7 promoter | T7 RNA polymerase promoter sequence | 1-17 |
| Cap | A modified 5’-cap1 structure ($m^7G$-5'-ppp-5'-$Am^{2'}$) | 18-19 |
| 5’-UTR | 5´-untranslated region derived from rat insulin2 mRNA (NM_019130.2) with an optimized Kozak sequence | 20-77 |
| Ccnd2 T280A | Sequence encoding rat Ccnd2 (NM_022267.2; in bold), with introduced mutation T280A (threonine to alanine at position 280); stop codons: 942-947 (underlined) | 78-947 |
| 3’-UTR | The 3´-untranslated region derived from rat insulin2 mRNA (NM_019130.2) | 948-1005 |
| poly(A) | A 110-nucleotide poly(A)-tail consisting of a stretch of 30 adenosine residues, followed by a 10-nucleotide linker sequence and another 70 adenosine residues. | 1006-1115 |

Sequence:

TAATACGACTCACTATAAGGAGCCCTAAGTGACCAGCTACAGTCGGAAAC 50<br>
CATCAGCAAGCAGGTCATTGTGCCAAC**ATGGAGCTGCTGTGCTGTGAGGT** 100<br>
**GGACCCGGTCCGCAGGGCCGTGCCGGACCGCAACCTGCTGGAAGACCGCG** 150<br>
**TCCTGCAGAACCTGTTGACTATCGAGGAGCGCTACCTCCCGCAGTGTTCC** 200<br>
**TATTTCAAGTGCGTGCAGAAGGACATCCAGCCGTACATGCGCAGGATGGT** 250<br>
**GGCTACCTGGATGCTAGAGGTCTGTGAGGAACAGAAGTGTGAAGAAGAGG** 300<br>
**TCTTTCCTCTGGCCATGAATTACCTGGACCGTTTCTTGGCTGGAGTCCCG** 350<br>
**ACTCCTAAGACCCATCTCCAGCTCCTGGGCGCTGTGTGCATGTTCCTAGC** 400<br>
**TTCCAAGCTGAAAGAGACCATCCCGCTGACTGCCGAAAAGCTGTGTATTT** 450<br>
**ACACCGACAATTCTGTGAAACCCCAGGAGCTGCTGGAGTGGGAACTGGTG** 500<br>
**GTGCTGGGTAAGCTGAAGTGGAACCTGGCTGCAGTAACCCCTCACGACTT** 550<br>
**CATTGAGCACATCCTACGCAAGCTGCCCCAGCAGAAGGAGAAGCTGTCCC** 600<br>
**TGATCCGCAAGCATGCGCAGACCTTCATCGCTCTGTGTGCTACCGACTTC** 650<br>
**AAGTTTGCCATGTACCCGCCATCGATGATCGCAACTGGAAGCGTGGGAGC** 700<br>
**AGCCATCTGCGGGCTTCAGCAGGACGAGGAAGTGAATGCACTCACGTGCG** 750<br>
**ATGCCCTGACGGAGCTGCTGGCCAAGATCACCCACACCGATGTGGATTGT** 800<br>
**CTCAAAGCCTGCCAGGAGCAAATCGAGGCTGTGCTGCTTAACAGCCTTCA** 850<br>
**GCAGTTCCGTCAAGAGCAGCACAACGGCTCCAAGTCTGTGGAAGATCCGG** 900<br>
**ACCAAGCCACCGCCCCTACAGACGTGCGGGATGTTGACCTG**TGATAAGCC 950<br>
CACCACTACCCTGTCCACCCCTCTGCAATGAATAAAACCTTTGAAAGAGC 1000<br>
ACTACAAAAAAAAAAAAAAAAAAAAAAAAAAAAAAGCATATGACTAAAAA 1050<br>
AAAAAAAAAAAAAAAAAAAAAAAAAAAAAAAAAAAAAAAAAAAAAAAAAA 1100<br>
AAAAAAAAAAAAAAA 1115

Table S6. DNA template sequence encoding rat Ccnd3 IVT mRNA.

| Element | Description | Position |
|---|---|---|
| T7 promoter | T7 RNA polymerase promoter sequence | 1-17 |
| Cap | A modified 5'-cap1 structure ($m^7G$-5'-ppp-5'-$Am^{2'}$) | 18-19 |
| 5'-UTR | 5´-untranslated region derived from rat insulin2 mRNA (NM_019130.2) with an optimized Kozak sequence | 20-77 |
| Ccnd3 | Sequence encoding rat Ccnd3 (NM_012766.3; in bold); stop codons: 954-959 (underlined) | 78-959 |
| 3'-UTR | The 3´-untranslated region derived from rat insulin2 mRNA (NM_019130.2) | 960-1017 |
| poly(A) | A 110-nucleotide poly(A)-tail consisting of a stretch of 30 adenosine residues, followed by a 10-nucleotide linker sequence and another 70 adenosine residues. | 1018-1127 |

Sequence:

TAATACGACTCACTATAAGGAGCCCTAAGTGACCAGCTACAGTCGGAAAC **50**<br>
CATCAGCAAGCAGGTCATTGTGCCAAC**ATGGAGCTGCTGTGTTGCGAGGG** **100**<br>
**CACCCGGCACGCGCCCCGGGCCGGGCCCGACCCGCGGCTACTGGGGACC** **150**<br>
**AGCGTGTCCTGCAGAGTTTGCTCCGCTTGGAGGAGCGCTACGTGCCGCGA** **200**<br>
**GCCTCCTACTTCCAGTGCGTGCAAAAGGAGATCAAGCCGCACATGCGGAA** **250**<br>
**GATGCTGGCGTACTGGATGCTGGAGGTGTGTGAGGAGCAGCGCTGCGAGG** **300**<br>
**AGGATGTCTTCCCTCTGGCTATGAACTACCTGGATCGCTACCTGTCCTGC** **350**<br>
**GTCCCCACCCGAAAGGCGCAACTGCAGCTTCTAGGTACCGTCTGCCTGTT** **400**<br>
**GCTGGCCTCCAAGCTGCGCGAAACCACGCCCCTGACTATTGAGAAGCTCT** **450**<br>
**GCATCTATACGGACCAAGCTGTGGCTCCCTGGCAGTTGCGGGAATGGGAG** **500**<br>
**GTGCTGGTCCTGGGGAAGCTCAAGTGGGACCTGGCTGCTGTGATTGCGCA** **550**<br>
**CGACTTCCTGGCCCTGATTCTGCACCGCCTCTCTCTGCCCAGTGACCGGC** **600**<br>
**AGGCACTGGTCAAAAAGCATGCTCAGACCTTTTTGGCCCTCTGTGCCACA** **650**<br>
**GATTACACCTTTGCGATGTACCCTCCATCCATGATCGCCACGGGCAGCAT** **700**<br>
**CGGGGCTGCAGTGCTAGGCCTGGGTGCCTGCTCTATGTCTGCAGATGAGC** **750**<br>
**TCACAGAGCTGCTGGCGGGAATCACAGGCACTGAAGTGGACTGCCTGCGT** **800**<br>
**GCCTGCCAGGAGCAGATCGAAGCTGCCCTCAGGGAGAGCCTCAGGGAAGC** **850**<br>
**TGCTCAGACAGCCCCCAGCCCCGTGCCCAAAGCCCCCCGGGGCTCTAGCA** **900**<br>
**GCCAGGGGCCCAGTCAGACCAGCACTCCCACAGATGTCACAGCCATCCAC** **950**<br>
**CTG**TAGTAAGCCCACCACTACCCTGTCCACCCCTCTGCAATGAATAAAAC **1000**<br>
CTTTGAAAGAGCACTACAAAAAAAAAAAAAAAAAAAAAAAAAAAAAAGCA **1050**<br>
TATGACTAAAAAAAAAAAAAAAAAAAAAAAAAAAAAAAAAAAAAAAAAAA **1100**<br>
AAAAAAAAAAAAAAAAAAAAAAAAAAA **1127**

Table S7. DNA template sequence encoding rat Ccnd3 T283A IVT mRNA.

| Element | Description | Position |
|---|---|---|
| T7 promoter | T7 RNA polymerase promoter sequence | 1-17 |
| Cap | A modified 5'-cap1 structure ($m^7G$-5'-ppp-5'-$Am^{2'}$) | 18-19 |
| 5'-UTR | 5´-untranslated region derived from rat insulin2 mRNA (NM_019130.2) with an optimized Kozak sequence | 20-77 |
| Ccnd3 T283A | Sequence encoding rat Ccnd3 (NM_012766.3; in bold), with introduced mutation T283A (threonine to alanine at position 283); stop codons: 954-959 (underlined) | 78-959 |
| 3'-UTR | The 3´-untranslated region derived from rat insulin2 mRNA (NM_019130.2) | 960-1017 |
| poly(A) | A 110-nucleotide poly(A)-tail consisting of a stretch of 30 adenosine residues, followed by a 10-nucleotide linker sequence and another 70 adenosine residues. | 1018-1127 |

Sequence:

TAATACGACTCACTATAAGGAGCCCTAAGTGACCAGCTACAGTCGGAAAC 50<br>
CATCAGCAAGCAGGTCATTGTGCCAAC**ATGGAGCTGCTGTGTTGCGAGGG** 100<br>
**CACCCGGCACGCGCCCCGGGCCGGGCCCGACCCGCGGCTACTGGGGGACC** 150<br>
**AGCGTGTCCTGCAGAGTTTGCTCCGCTTGGAGGAGCGCTACGTGCCGCGA** 200<br>
**GCCTCCTACTTCCAGTGCGTGCAAAAGGAGATCAAGCCGCACATGCGGAA** 250<br>
**GATGCTGGCGTACTGGATGCTGGAGGTGTGTGAGGAGCAGCGCTGCGAGG** 300<br>
**AGGATGTCTTCCCTCTGGCTATGAACTACCTGGATCGCTACCTGTCCTGC** 350<br>
**GTCCCCACCCGAAAGGCGCAACTGCAGCTTCTAGGTACCGTCTGCCTGTT** 400<br>
**GCTGGCCTCCAAGCTGCGCGAAACCACGCCCCTGACTATTGAGAAGCTCT** 450<br>
**GCATCTATACGGACCAAGCTGTGGCTCCCTGGCAGTTGCGGGAATGGGAG** 500<br>
**GTGCTGGTCCTGGGGAAGCTCAAGTGGGACCTGGCTGCTGTGATTGCGCA** 550<br>
**CGACTTCCTGGCCCTGATTCTGCACCGCCTCTCTCTGCCCAGTGACCGGC** 600<br>
**AGGCACTGGTCAAAAAGCATGCTCAGACCTTTTTGGCCCTCTGTGCCACA** 650<br>
**GATTACACCTTTGCGATGTACCCTCCATCCATGATCGCCACGGGCAGCAT** 700<br>
**CGGGGCTGCAGTGCTAGGCCTGGGTGCCTGCTCTATGTCTGCAGATGAGC** 750<br>
**TCACAGAGCTGCTGGCGGGAATCACAGGCACTGAAGTGGACTGCCTGCGT** 800<br>
**GCCTGCCAGGAGCAGATCGAAGCTGCCCTCAGGGAGAGCCTCAGGGAAGC** 850<br>
**TGCTCAGACAGCCCCCAGCCCCGTGCCCAAAGCCCCCCGGGGCTCTAGCA** 900<br>
**GCCAGGGGCCCAGTCAGACCAGCGCTCCCACAGATGTCACAGCCATCCAC** 950<br>
**CTG**<u>TAGTAA</u>GCCCACCACTACCCTGTCCACCCCTCTGCAATGAATAAAAC 1000<br>
CTTTGAAAGAGCACTACAAAAAAAAAAAAAAAAAAAAAAAAAAAAAAGCA 1050<br>
TATGACTAAAAAAAAAAAAAAAAAAAAAAAAAAAAAAAAAAAAAAAAAAA 1100<br>
AAAAAAAAAAAAAAAAAAAAAAAAAAA 1127

Table S8. DNA template sequence encoding rat Cdk4 IVT mRNA.

| Element | Description | Position |
|---|---|---|
| T7 promoter | T7 RNA polymerase promoter sequence | 1-17 |
| Cap | A modified 5'-cap1 structure (m$^7$G-5'-ppp-5'-Am$^{2'}$) | 18-19 |
| 5'-UTR | 5´-untranslated region derived from rat insulin2 mRNA (NM_019130.2) with an optimized Kozak sequence | 20-77 |
| Cdk4 | Sequence encoding rat Cdk4 (NM_053593.3; in bold); stop codons: 987-992 (underlined) | 78-992 |
| 3'-UTR | The 3´-untranslated region derived from rat insulin2 mRNA (NM_019130.2) | 993-1050 |
| poly(A) | A 110-nucleotide poly(A)-tail consisting of a stretch of 30 adenosine residues, followed by a 10-nucleotide linker sequence and another 70 adenosine residues. | 1051-1160 |

Sequence:

TAATACGACTCACTATAAGGAGCCCTAAGTGACCAGCTACAGTCGGAAAC 50
CATCAGCAAGCAGGTCATTGTGCCAAC**ATGGCTACCACTCGATATGAACC** 100
**CGTGGCTGAAATTGGTGTCGGTGCCTATGGACGGTGTACAAAGCCCGAG** 150
**ATCCCCACAGTGGCCACTTTGTGGCTCTCAAGAGTGTGAGAGTTCCTAAT** 200
**GGAGGAGCAGCTGGAGGGGGCCTTCCCGTCAGCACAGTTCGTGAGGTGGC** 250
**CTTGTTAAGAAGGCTGGAGGCCTTTGAACATCCCAATGTTGTACGGCTGA** 300
**TGGATGTCTGTGCTACTTCCCGAACTGATCGGGACATCAAGGTCACCTTA** 350
**GTGTTTGAGCATATAGACCAGGACCTACGGACATACCTGGACAAAGCACC** 400
**TCCGCCGGGCTTGCCTGTTGAGACCATTAAGGATCTGATGCGCCAGTTTC** 450
**TAAGCGGCCTAGATTTCCTTCATGCAAACTGCATTGTTCACCGGGACCTG** 500
**AAGCCAGAGAACATTCTAGTGACAAGTAATGGGACAGTTAAGCTGGCCGA** 550
**CTTTGGCCTAGCCAGAATCTACAGCTACCAGATGGCCCTCACGCCTGTGG** 600
**TTGTTACGCTCTGGTACCGGGCTCCTGAAGTTCTTCTGCAGTCTACATAT** 650
**GCAACGCCTGTGGATATGTGGAGTGTTGGCTGTATCTTCGCAGAGATGTT** 700
**TCGCCGGAAGCCTCTCTTCTGTGGGAACTCTGAGGCTGACCAGCTGGGCA** 750
**AAATCTTTGATCTCATTGGATTGCCTCCAGAAGACGACTGGCCTCGAGAG** 800
**GTCTCTCTTCCTCGAGGAGCCTTTTCCCCCAGAGGACCTCGGCCAGTGCA** 850
**GTCAGTGGTGCCGGAGATGGAGGAATCTGGAGCGCAGTTGCTGCTGGAAA** 900
**TGCTGACCTTTAATCCACTTAAGCGAATCTCTGCCTTCCGAGCCCTGCAG** 950
**CACTCTTACCTGCACAAGGAGGAAAGTGACCCGGAG**TGATAAGCCCACCA 1000
CTACCCTGTCCACCCCTCTGCAATGAATAAAACCTTTGAAAGAGCACTAC 1050
AAAAAAAAAAAAAAAAAAAAAAAAAAAAAAGCATATGACTAAAAAAAAAA 1100
AAAAAAAAAAAAAAAAAAAAAAAAAAAAAAAAAAAAAAAAAAAAAAAAAA 1150
AAAAAAAAAA 1160

Table S9. DNA template sequence encoding rat Cdk6 IVT mRNA.

| Element | Description | Position |
|---|---|---|
| T7 promoter | T7 RNA polymerase promoter sequence | 1-17 |
| Cap | A modified 5'-cap1 structure ($m^7G$-5'-ppp-5'-$Am^{2'}$) | 18-19 |
| 5'-UTR | 5´-untranslated region derived from rat insulin2 mRNA (NM_019130.2) with an optimized Kozak sequence | 20-77 |
| Cdk6 | Sequence encoding rat Cdk6 (NM_001191861.3; in bold); stop codons: 1056-1061 (underlined) | 78-1061 |
| 3'-UTR | The 3´-untranslated region derived from rat insulin2 mRNA (NM_019130.2) | 1062-1118 |
| poly(A) | A 110-nucleotide poly(A)-tail consisting of a stretch of 30 adenosine residues, followed by a 10-nucleotide linker sequence and another 70 adenosine residues. | 1119-1229 |

Sequence:

TAATACGACTCACTATAAGGAGCCCTAAGTGACCAGCTACAGTCGGAAAC 50
CATCAGCAAGCAGGTCATTGTGCCAAC**ATGGAGAAGGACAGCCTGAGTCG** 100
**CGCCGACCAGCAGTATGAGTGCGTGGCGGAGATCGGGGAAGGCGCCTACG** 150
**GGAAGGTGTTCAAGGCCCGCGACCTGAAGAACGGCGGCCGCTTCGTGGCT** 200
**CTGAAGCGCGTGCGAGTGCAGACCGGAGAGGAGGGCATGCCGCTCTCCAC** 250
**CATCCGCGAGGTGGCGGTGCTGAGGCACCTGGAGACCTTTGAGCACCCCA** 300
**ACGTGGTCAGGTTGTTTGACGTGTGCACAGTGTCACGGACAGACAGAGAA** 350
**ACTAAACTTACGCTAGTGTTTGAGCATGTTGATCAAGACTTGACCACTTA** 400
**CTTGGATAAAGTTCCAGAACCCGGTGTGCCCACAGAGACCATAAAGGATA** 450
**TGATGTTTCAGCTTCTCCGAGGTCTGGACTTCCTCCATTCTCACAGAGTA** 500
**GTGCATCGTGACCTGAAGCCACAGAACATTCTGGTGACCAGCAGTGGACA** 550
**AATAAAACTGGCTGACTTCGGCCTTGCCCGCATCTACAGTTTTCAGATGG** 600
**CCCTTACCTCGGTGGTCGTCACGCTGTGGTACCGAGCCCCGGAAGTCCTG** 650
**CTCCAGTCCAGCTACGCCACCCCCGTGGACCTCTGGAGTGTTGGCTGCAT** 700
**CTTTGCAGAACTGTTTCGCAGAAAGCCTCTTTTTCGTGGAAGTTCAGACG** 750
**TGGATCAACTAGGGAAAATCTTGGACGTCATCGGACTCCCAGGAGAAGAA** 800
**GACTGGCCTAGGGATGTTGCTCTTCCCAGACAGGCTTTTCACTCCAAATC** 850
**TGCCCAACCCATCGAGAAGTTTGTGACAGACATCGACGAGCTAGGCAAAG** 900
**ACCTCCTTCTGAAATGCTTGACGTTTAATCCAGCTAAAAGAATATCCGCT** 950
**TATGGCGCCCTGAATCACCCGTACTTCCAAGACCTGGAGAGATACAAGGA** 1000
**CAACCTGCATTCTCACCTGTCGTCCAGCCAGAGCACCTCGGAGCTGAACA** 1050
**CAGCC**TGATAAGCCCACCACTACCCTGTCCACCCCTCTGCAATGAATAAA 1100
ACCTTTGAAAGAGCACTACAAAAAAAAAAAAAAAAAAAAAAAAAAAAAAG 1150
CATATGACTAAAAAAAAAAAAAAAAAAAAAAAAAAAAAAAAAAAAAAAAA 1200
AAAAAAAAAAAAAAAAAAAAAAAAAAAAA 1229

Table S10. DNA template sequence encoding human CCND1 T286A IVT mRNA.

| Element | Description | Position |
|---|---|---|
| T7 promoter | T7 RNA polymerase promoter sequence | 1-17 |
| Cap | A modified 5’-cap1 structure ($m^7G$-5'-ppp-5'-$Am^{2'}$) | 18-19 |
| 5’-UTR | 5´-untranslated region derived from human insulin mRNA (NM_000207.3) | 20-79 |
| CCND1 T286A | Sequence encoding human CCND1 (NM_053056.3; in bold), with introduced mutation T286A (threonine to alanine at position 286); stop codons: 965-970 (underlined) | 80-970 |
| 3’-UTR | The 3´-untranslated region derived from human insulin mRNA (NM_000207.3) | 971-1043 |
| poly(A) | A 110-nucleotide poly(A)-tail consisting of a stretch of 30 adenosine residues, followed by a 10-nucleotide linker sequence and another 70 adenosine residues. | 1044-1153 |

Sequence:

| | |
|---|---|
| TAATACGACTCACTATAAGGAGCCCTCCAGGACAGGCTGCATCAGAAGAG | 50 |
| GCCATCAAGCAGATCACTGTCCTTCTGCC**ATGGAACACCAGCTCCTGTGC** | 100 |
| **TGCGAAGTGGAAACCATCCGCCGCGCGTACCCCGATGCCAACCTCCTCAA** | 150 |
| **CGACCGGGTGCTGCGGGCCATGCTGAAGGCGGAGGAGACCTGCGCGCCCT** | 200 |
| **CGGTGTCCTACTTCAAATGTGTGCAGAAGGAGGTCCTGCCGTCCATGCGG** | 250 |
| **AAGATCGTCGCCACCTGGATGCTGGAGGTCTGCGAGGAACAGAAGTGCGA** | 300 |
| **GGAGGAGGTCTTCCCGCTGGCCATGAACTACCTGGACCGCTTCCTGTCGC** | 350 |
| **TGGAGCCCGTGAAAAAGAGCCGCCTGCAGCTGCTGGGGGCCACTTGCATG** | 400 |
| **TTCGTGGCCTCTAAGATGAAGGAGACCATCCCCCTGACGGCCGAGAAGCT** | 450 |
| **GTGCATCTACACCGACAACTCCATCCGGCCCGAGGAGCTGCTGCAAATGG** | 500 |
| **AGCTGCTCCTGGTGAACAAGCTCAAGTGGAACCTGGCCGCAATGACCCCG** | 550 |
| **CACGATTTCATTGAACACTTCCTCTCCAAAATGCCAGAGGCGGAGGAGAA** | 600 |
| **CAAACAGATCATCCGCAAACACGCGCAGACCTTCGTTGCCCTCTGTGCCA** | 650 |
| **CAGATGTGAAGTTCATTTCCAATCCGCCCTCCATGGTGGCAGCGGGGAGC** | 700 |
| **GTGGTGGCCGCAGTGCAAGGCCTGAACCTGAGGAGCCCCAACAACTTCCT** | 750 |
| **GTCCTACTACCGCCTCACACGCTTCCTCTCCAGAGTGATCAAGTGTGACC** | 800 |
| **CGGACTGCCTCCGGGCCTGCCAGGAGCAGATCGAAGCCCTGCTGGAGTCA** | 850 |
| **AGCCTGCGCCAGGCCCAGCAGAACATGGACCCCAAGGCCGCCGAGGAGGA** | 900 |
| **GGAAGAGGAGGAAGAGGAAGTGGACCTGGCTTGCGCACCCACCGACGTGC** | 950 |
| **GGGACGTGGACATC**TGATAAACGCAGCCCGCAGGCAGCCCCACACCCGCC | 1000 |
| GCCTCCTGCACCGAGAGAGATGGAATAAAGCCCTTGAACCAGCAAAAAAA | 1050 |
| AAAAAAAAAAAAAAAAAAAAAAGCATATGACTAAAAAAAAAAAAAAAAAA | 1100 |
| AAAAAAAAAAAAAAAAAAAAAAAAAAAAAAAAAAAAAAAAAAAAAAAAAA | 1150 |
| AAA | 1153 |

Table S11. DNA template sequence encoding human CDK4 IVT mRNA.

| Element | Description | Position |
|---|---|---|
| T7 promoter | T7 RNA polymerase promoter sequence | 1-17 |
| Cap | A modified 5'-cap1 structure ($m^7G$-5'-ppp-5'-$Am^{2'}$) | 18-19 |
| 5'-UTR | 5´-untranslated region derived from human insulin mRNA (NM_000207.3) | 20-79 |
| CDK4 | Sequence encoding human CDK4 (NM_000075.4; in bold); stop codons: 989-994 (underlined) | 80-994 |
| 3'-UTR | The 3´-untranslated region derived from human insulin mRNA (NM_000207.3) | 995-1067 |
| poly(A) | A 110-nucleotide poly(A)-tail consisting of a stretch of 30 adenosine residues, followed by a 10-nucleotide linker sequence and another 70 adenosine residues. | 1068-1177 |

Sequence:

TAATACGACTCACTATAAGGAGCCCTCCAGGACAGGCTGCATCAGAAGAG 50<br>
GCCATCAAGCAGATCACTGTCCTTCTGCC**ATGGCTACCTCTCGATATGAG** 100<br>
**CCAGTGGCTGAAATTGGTGTCGGTGCCTATGGGACAGTGTACAAGGCCCG** 150<br>
**TGATCCCCACAGTGGCCACTTTGTGGCCCTCAAGAGTGTGAGAGTCCCCA** 200<br>
**ATGGAGGAGGAGGTGGAGGAGGCCTTCCCATCAGCACAGTTCGTGAGGTG** 250<br>
**GCTTTACTGAGGCGACTGGAGGCTTTTGAGCATCCCAATGTTGTCCGGCT** 300<br>
**GATGGACGTCTGTGCCACATCCCGAACTGACCGGGAGATCAAGGTAACCC** 350<br>
**TGGTGTTTGAGCATGTAGACCAGGACCTAAGGACATATCTGGACAAGGCA** 400<br>
**CCCCCACCAGGCTTGCCAGCCGAAACGATCAAGGATCTGATGCGCCAGTT** 450<br>
**TCTAAGAGGCCTAGATTTCCTTCATGCCAATTGCATCGTTCACCGAGATC** 500<br>
**TGAAGCCAGAGAACATTCTGGTGACAAGTGGTGGAACAGTCAAGCTGGCT** 550<br>
**GACTTTGGCCTGGCCAGAATCTACAGCTACCAGATGGCACTTACACCCGT** 600<br>
**GGTTGTTACACTCTGGTACCGAGCTCCCGAAGTTCTTCTGCAGTCCACAT** 650<br>
**ATGCAACACCTGTGGACATGTGGAGTGTTGGCTGTATCTTTGCAGAGATG** 700<br>
**TTTCGTCGAAAGCCTCTCTTCTGTGGAAACTCTGAAGCCGACCAGTTGGG** 750<br>
**CAAAATCTTTGACCTGATTGGGCTGCCTCCAGAGGATGACTGGCCTCGAG** 800<br>
**ATGTATCCCTGCCCCGTGGAGCCTTTCCCCCAGAGGGCCCCGCCCAGTG** 850<br>
**CAGTCGGTGGTACCTGAGATGGAGGAGTCGGGAGCACAGCTGCTGCTGGA** 900<br>
**AATGCTGACTTTTAACCCACACAAGCGAATCTCTGCCTTTCGAGCTCTGC** 950<br>
**AGCACTCTTATCTACATAAGGATGAAGGTAATCCGGAG**TGATAAACGCAG 1000<br>
CCCGCAGGCAGCCCCACACCCGCCGCCTCCTGCACCGAGAGAGATGGAAT 1050<br>
AAAGCCCTTGAACCAGCAAAAAAAAAAAAAAAAAAAAAAAAAAAAAAGCA 1100<br>
TATGACTAAAAAAAAAAAAAAAAAAAAAAAAAAAAAAAAAAAAAAAAAAA 1150<br>
AAAAAAAAAAAAAAAAAAAAAAAAAAA 1177

Table S12. DNA template sequence encoding human MYC IVT mRNA.

| Element | Description | Position |
|---|---|---|
| T7 promoter | T7 RNA polymerase promoter sequence | 1-17 |
| Cap | A modified 5’-cap1 structure ($m^7G$-5'-ppp-5'-$Am^{2'}$) | 18-19 |
| 5’-UTR | 5´-untranslated region derived from human insulin mRNA (NM_000207.3) | 20-79 |
| MYC | Sequence encoding human MYC (NM_002467.6; in bold); stop codons: 1397-1402 (underlined) | 80-1402 |
| 3’-UTR | The 3´-untranslated region derived from human insulin mRNA (NM_000207.3) | 1403-1475 |
| poly(A) | A 110-nucleotide poly(A)-tail consisting of a stretch of 30 adenosine residues, followed by a 10-nucleotide linker sequence and another 70 adenosine residues. | 1476-1585 |

Sequence:

```
TAATACGACTCACTATAAGGAGCCCTCCAGGACAGGCTGCATCAGAAGAG    50
GCCATCAAGCAGATCACTGTCCTTCTGCCATGCCCCTCAACGTTAGCTTC   100
ACCAACAGGAACTATGACCTCGACTACGACTCGGTGCAGCCGTATTTCTA   150
CTGCGACGAGGAGGAGAACTTCTACCAGCAGCAGCAGCAGAGCGAGCTGC   200
AGCCCCCGGCGCCCAGCGAGGATATCTGGAAGAAATTCGAGCTGCTGCCC   250
ACCCCGCCCCTGTCCCCTAGCCGCCGCTCCGGGCTCTGCTCGCCCTCCTA   300
CGTTGCGGTCACACCCTTCTCCCTTCGGGGAGACAACGACGGCGGTGGCG   350
GGAGCTTCTCCACGGCCGACCAGCTGGAGATGGTGACCGAGCTGCTGGGA   400
GGAGACATGGTGAACCAGAGTTTCATCTGCGACCCGGACGACGAGACCTT   450
CATCAAAAACATCATCATCCAGGACTGTATGTGGAGCGGCTTCTCGGCCG   500
CCGCCAAGCTCGTCTCAGAGAAGCTGGCCTCCTACCAGGCTGCGCGCAAA   550
GACAGCGGCAGCCCGAACCCCGCCCGCGGCCACAGCGTCTGCTCCACCTC   600
CAGCTTGTACCTGCAGGATCTGAGCGCCGCCGCCTCAGAGTGCATCGACC   650
CCTCGGTGGTCTTCCCCTACCCTCTCAACGACAGCAGCTCGCCCAAGTCC   700
TGCGCCTCGCAAGACTCCAGCGCCTTCTCTCCGTCCTCGGATTCTCTGCT   750
CTCCTCGACGGAGTCCTCCCCGCAGGGCAGCCCCGAGCCCCTGGTGCTCC   800
ATGAGGAGACACCGCCCACCACCAGCAGCGACTCTGAGGAGGAACAAGAA   850
GATGAGGAAGAAATCGATGTTGTTTCTGTGGAAAAGAGGCAGGCTCCTGG   900
CAAAAGGTCAGAGTCTGGATCACCTTCTGCTGGAGGCCACAGCAAACCTC   950
CTCACAGCCCACTGGTCCTCAAGAGGTGCCACGTCTCCACACATCAGCAC  1000
AACTACGCAGCGCCTCCCTCCACTCGGAAGGACTATCCTGCTGCCAAGAG  1050
GGTCAAGTTGGACAGTGTCAGAGTCCTGAGACAGATCAGCAACAACCGAA  1100
AATGCACCAGCCCCAGGTCCTCGGACACCGAGGAGAATGTCAAGAGGCGA  1150
```

**ACACACAACGTCTTGGAGCGCCAGAGGAGGAACGAGCTAAAACGGAGCTT** 1200
**TTTTGCCCTGCGTGACCAGATCCCGGAGTTGGAAAACAATGAAAAGGCCC** 1250
**CCAAGGTAGTTATCCTTAAAAAAGCCACAGCATACATCCTGTCCGTCCAA** 1300
**GCAGAGGAGCAAAAGCTCATTTCTGAAGAGGACTTGTTGCGGAAACGACG** 1350
**AGAACAGTTGAAACACAAACTTGAACAGCTACGGAACTCTTGTGCG**TAAT 1400
AGACGCAGCCCGCAGGCAGCCCCACACCCGCCGCCTCCTGCACCGAGAGA 1450
GATGGAATAAAGCCCTTGAACCAGCAAAAAAAAAAAAAAAAAAAAAAAAA 1500
AAAAAGCATATGACTAAAAAAAAAAAAAAAAAAAAAAAAAAAAAAAAAAA 1550
AAAAAAAAAAAAAAAAAAAAAAAAAAAAAAAAAAA 1585

Table S13. List of antibodies used in the study

| Antibody | Vendor | Cat. No. | Clone | Dilution |
|---|---|---|---|---|
| Rabbit anti-Cyclin A2, IgG | Abcam | ab181591 | EPR17351 | 1:600 (IF) |
| Rabbit anti-Insulin, IgG | Abcam | ab181547 | EPR17359 | 1:400 (IF)<br>1:200 (FC) |
| Rabbit anti-Ki67 | Abcam | ab15580 | Polyclonal | 1:600 (IF) |
| Mouse anti-Cyclin D3, IgG | BD | 610279 | 1/Cyclin D3 | 1:200 (IF)<br>1:3000 (WB) |
| Goat anti-Rabbit HRP | Biorad | 1705046 | N/A | 1:20000 (WB) |
| Rabbit anti-Cyclin D2, IgG | CST | 3741 | D52F9 | 1:100 (IF) |
| Rabbit anti-Phospho-Histone H3 (Ser10), IgG | CST | 3377 | D2C8 | 1:600 (IF) |
| Rabbit anti-Phospho-Rb (Ser807/811), IgG | CST | 8516 | D20B12 | 1:600 (IF) |
| Mouse anti-Insulin, IgG | Exbio | 11-246-C100 | IN-05 | 1:200 (IF) |
| Rabbit anti-Cyclin D1 | NOVUS | NBP2-16054 | Polyclonal | 1:200 (IF)<br>1:2000 (WB) |
| Rabbit anti-CDK4 | NOVUS | NBP1-31308 | Polyclonal | 1:200 (IF);<br>1:2000 (WB) |
| Rabbit anti-Cyclin D2 | Proteintech | 10934-1-AP | Polyclonal | 1:2000 (WB) |
| Mouse anti-CDK6, IgG | Proteintech | 66278-1-Ig | 4B9C11 | 1:200 (IF)<br>1:8000 (WB) |
| Mouse anti-Beta-Actin, IgG | SigmaAldrich | A5441 | AC-15 | 1:5000 (WB) |
| Mouse anti-Insulin, IgG | SigmaAldrich | I2018 | K36AC10 | 1:100 (IF) |
| Donkey anti-Rabbit, IgG, AlexaFluor 488 | ThermoFisher | A32790 | N/A | 1:800 (IF) |
| Donkey anti-Mouse, IgG, AlexaFluor 555 | ThermoFisher | A31570 | N/A | 1:400 (IF) |
| Donkey anti-Rabbit, IgG, AlexaFluor 555 | ThermoFisher | A31572 | N/A | 1:400 (IF)<br>1:100 (FC) |
| Donkey anti-Mouse, IgG, AlexaFluor 647 | ThermoFisher | A31571 | N/A | 1:400 (IF) |
| Donkey anti-Rabbit, IgG, AlexaFluor Plus 647 | ThermoFisher | A32795 | N/A | 1:800 (IF) |
| Rabbit anti-Mouse HRP | ThermoFisher | 81-6720 | Polyclonal | 1:20000 (WB) |

Table S14. Sequences of primers used for RT-qPCR

| **Gene** | **Forward (5'→3')** | **Reverse (5'→3')** |
|---|---|---|
| *Ins1* | CCCTAAGTGACCAGCTACAATCAT | CGGGACTTGGGTGTGTAGAAG |
| *Ins2* | AAGTGACCAGCTACAGTCGG | ACCTCCAGTGCCAAGGTCTG |
| *Gcg* | AGACCGTTTACATCGTGGCT | GGCAATGTTGTTCCGGTTCC |
| *Sst* | CCCCAGACTCCGTCAGTTTC | CGCAGGGTCTAGTTGAGCAT |
| *Gck* | TGCCGAGATGCTCTTTGACTACA | GGGTTCGCTGAGCTTTCATCC |
| *Ucn3* | AGAGCAAAGTCCTCTTACAGGGA | CCCCGGTCGTTTTTGACCTT |
| *Pdx1* | TTCCCGAATGGAACCGAGACT | TCCACTTCATGCGACGGTTT |
| *Mafa* | GCCCCGAGAACGGTGAATAC | AGGGAGTTCCTCCGGGTTTT |
| *Neurod1* | GGCTCCAGGGTTATGAGATCG | GCCAAGCGCAGTGTCTCTAT |
| *Nkx6-1* | AGAGAGCACGCTTGGCCTATT | AATAGTAAAGCCGGGCGAGCA |
| *Nkx2-2* | ACCGAGGGCCTCCAATACTC | GGCACGTTTCATCTTGTAGCG |
| *Pax4* | CAATCGAGTCCTTCGGGCAC | GCCCACGCTGGAACTCTTTC |
| *Ccnd1* | CAGATCATCCGCAAACATGCACA | CAGATCATCCGCAAACATGCACA |
| *Ccnd2* | TCACCCCTCACGACTTCATTGAG | ATGACGAACACGCCTCTCTCTTG |
| *Ccnd3* | CTCAGACCTTTTTGGCCCTCTGT | CTGAGGGGGCCTGTCCCAAACTA |
| *Ccne1* | CCGTCTTGAATTGGGGCAATAG | TGGTGCAACTTTGGAGGGTAGAT |
| *Ccne2* | TCAGGAGACGTTCATCCAAATAGC | AAAAGGCACCATCCAGTCTACAC |
| *Ccna1* | CTGACCGTTCCAACCACCAACCA | GCTCACTCAGGCAAGGCACAATC |
| *Ccna2* | TAAACAGCCTGCCTTCACCATTC | ACTTCAACTAGCCAGTCCACAAG |
| *Ccnb1* | GGATGTGCACCTGCCGAAGAATA | ACAACTGTCCTGCATGAACCGAT |
| *Cdk1* | AGTACGGCAATCCGGGAAATCTC | ATGAACTGGCCAGGAGGGATAGA |
| *Cdk2* | AGCCTTTGGAGTCCCTGTCCGTA | CCGAGCCCACTTGGGGAAACTTG |
| *Cdk4* | GTCTATGGTCTGGCCCGAAGCG | TACACCGTCCCATAGGCACCGA |
| *Cdk6* | AATCTGCCCAACCCATCGAGAA | GGTCCTCAGCCAGCAGTCTGTAA |
| *Foxm1* | ATCCAGTGGCTTGGAAAGATGAG | GCCTGGCTTGGCAATGTGCTTAA |
| *Aurkb* | CAGCTCCGCCGAGAGATCGAAAT | TGCATCCGCCCTTCAATCATCTC |
| *Cdkn2a* | GTCGTGCGGTATTTGCGGTATC | TCTCGCGTTGCCAGAAGTGAAG |
| *Cdkn2c* | TTCTGCGAGACGGATGGAAAGTG | CCGGATTCCCAAGTTTCATAACCT |

| | | |
|---|---|---|
| *Cdkn2d* | CACCGGTAGCCACTATGCTTCTG | AGAGCAACTGCTGGACTTCCAAA |
| *Cdkn1c* | AATGCGAACGGTGCGATCAAGA | TCCGGTTCCTGCTACATGAATGA |
| *Actb* | TATCGGCAATGAGCGGTTCC | AAAACGCAGCTCAGTAACAGTCC |
| *Hprt* | ATGACCAGTCAACGGGGGACATA | CCAACAAAGTCTGGCCTGTATCC |
